# ERF and WRKY transcription factors regulate *IDA* and abscission timing in Arabidopsis

**DOI:** 10.1101/2023.09.12.557497

**Authors:** Sergio Galindo-Trigo, Anne-Maarit Bågman, Takashi Ishida, Shinichiro Sawa, Siobhán M. Brady, Melinka A. Butenko

## Abstract

Plants shed organs like leaves, petals or fruits through the process of abscission. Monitoring cues like age, resource availability, biotic and abiotic stresses allows plants to abscise organs in a timely manner. How these signals are integrated in the molecular pathways that drive abscission is largely unknown. The *INFLORESCENCE DEFICIENT IN ABSCISSION* (*IDA*) gene is one of the main drivers of floral organ abscission in Arabidopsis and is known to transcriptionally respond to most abscission-regulating cues. Interrogating the *IDA* promoter *in silico* and *in vitro* we identified transcription factors that can potentially modulate *IDA* expression. We functionally characterized the importance of ERF and WRKY binding sites for *IDA* expression during floral organ abscission, with WRKYs being of special relevance to mediate *IDA* upregulation in response to biotic stress in tissues destined for separation. We further characterized WRKY57 as a positive regulator of *IDA* and *IDA*-*like* gene expression in abscission zones. Our findings highlight the promise of promoter element-targeted approaches to modulate the responsiveness of the IDA signaling pathway to harness controlled abscission timing for improved crop productivity.

**Highlight:** ERF and WRKY transcription factors distinctly contribute to the regulation of *IDA* expression and thereby abscission timing. WRKY57 modulates abscission via redundant IDA/IDA-like peptides.

## Introduction

Abscission is the developmentally programmed process of cell separation. Indeterminate growth and a modular developmental plan allow plants to shed organs that are no longer needed. Abscission can take place in leaf petioles, floral organs, flower pedicels, fruits or seeds, to name a few. The ubiquitous presence of abscission across plant organs and developmental phases provides these sessile organisms with flexibility to prioritize resource allocation and a very effective strategy to minimize disease spread. On the other hand, untimely or uncontrolled abscission has profound negative consequences for agriculture. Indeed, as humankind has domesticated plants, seed abscission has been selected against in crops, hindering a process that naturally evolved to aid seed dispersal in benefit of more efficient and plentiful harvests (Pickersgill, 2007). Controlled abscission is still actively sought-after in breeding programs. Understanding the complex set of cues that influence abscission occurrence or its timing is thus highly relevant.

There is a broad spectrum of cues that influence abscission. The phytohormones auxin and ethylene have antagonistic effects on abscission (Addicott et al., 1955; Abeles and Rubinstein, 1964; Louie Jr and Addicott, 1970; Elmo M. Beyer and Morgan, 1971; Meir et al., 2006; Meir et al., 2010; Basu et al., 2013; Meir et al., 2015). The generally accepted model suggests the competence to abscise is blocked by auxin efflux from the organ into the abscission zone (AZ). Ethylene is a positive effector of abscission and ethylene sensitivity acquisition is a milestone for abscission induction (Abeles and Rubinstein, 1964; Burg, 1968; Reid, 1985; Brown, 1997; Cin et al., 2005; Merelo et al., 2017; Botton and Ruperti, 2019). Carbohydrate availability is a well-known factor regulating abscission induction (Sawicki et al., 2015). Carbohydrate starvation triggers abscission, as documented in multiple plant species after shading or defoliation (Addicott et al., 1955; Aloni et al., 1997; Peng et al., 2013). Exogenous cues like water availability and pathogens also regulate abscission induction (Patharkar and Walker, 2016; Reichardt et al., 2020). Plants sense infections in leaves and trigger abscission to diminish the spread of the disease (Ketring and Melouk, 1982; Ben-David et al., 1986; Glick et al., 2009; Scalschi et al., 2014). Other phytohormones (cytokinin, salicylic acid, jasmonic acid), developmental cues (senescence, fruit and seed development), as well as exogenous cues (light and temperature) are known to influence abscission (for review, see (Ma et al., 2021)). Understanding how the molecular pathways that execute abscission integrate these complex signals would greatly inform abscission-related breeding programs.

The abscission of floral organs in *Arabidopsis thaliana* (Arabidopsis) is the best characterized abscission model. AZs develop at the base of each floral organ (sepals, petals, and stamens) as two adjacent but distinct cell layers: the residuum and secession layers. Secession cells are located proximal to the abscising organ and form a lignified structure called lignin brace. The lignin brace is thought to focus cell wall degrading enzyme activity and provide local rigidity to facilitate shedding (Lee et al., 2018). Residuum cells make up the cell layer that remains on the flower receptacle after abscission occurs, differentiating into cuticle-bearing epidermal-like cells. Several transcription factors (TFs) are necessary for AZs to differentiate during flower development, including BLADE ON PETIOLE 1 (BOP1) and BOP2, ARABIDOPSIS THALIANA HOMEOBOX GENE 1 (ATH1), KNOTTED-LIKE FROM ARABIDOPSIS THALIANA 2 (KNAT2) and KNAT6 (McKim et al., 2008; Crick et al., 2022). When abscission zones have developed and floral organs are no longer required, abscission zones secrete the peptide INFLORESCENCE DEFICIENT IN ABSCISSION (IDA) to trigger the cell separation (Butenko et al., 2003). IDA is perceived in the AZ cells by leucine-rich repeat (LRR) receptor kinases (RKs) HAESA (HAE) and HAESA-LIKE 2 (HSL2) and their co-receptors, members of the family of somatic embryogenesis receptor kinases (SERKs; (Jinn et al., 2000; Cho et al., 2008; Stenvik et al., 2008; Meng et al., 2016; Santiago et al., 2016)). When the HAE-SERK/HSL2-SERK receptor complexes activate, they trigger an intracellular signaling cascade of MITOGEN-ACTIVATED PROTEIN KINASES (MAPKs; (Cho et al., 2008; Zhu et al., 2019)). The MAPK cascade inhibits negative transcriptional regulators of abscission, KNAT1 and AGAMOUS-LIKE 15, thereby allowing the progression of abscission (Wang et al., 2006; Shi et al., 2011; Butenko et al., 2012; Patharkar and Walker, 2015). Other regulators influence abscission indirectly by modulating the IDA-induced signaling pathway (Liljegren et al., 2009; Leslie et al., 2010; Burr et al., 2011; Liu et al., 2013; Gubert and Liljegren, 2014; Baer et al., 2016; Taylor et al., 2019)

The signaling cascade induced by IDA and HAE/HSL2 in AZs is a requisite for floral organ abscission. Double mutants *hae hsl2* retain all floral organs across floral positions in the inflorescence (Cho et al., 2008; Stenvik et al., 2008). Meanwhile, *ida* knockouts display a weaker abscission phenotype, with floral organs being loosely attached in floral positions in which siliques are elongating (Stenvik et al., 2008; Alling and Galindo-Trigo, 2023). The weaker abscission phenotype in *ida* mutants is likely due to functional redundancy with related IDA-like (IDL) peptides in AZs (Stenvik et al., 2008; Vie et al., 2015). A recent revisitation of the literature has proposed the IDA pathway could be responsible for the very last steps of separation mediating an increase in turgidity and cell expansion, while abscission activation and initiation of cell wall degradation would be mostly dependent on ethylene signaling (Meir et al., 2019). In this scenario, IDA would be one of several responses that ethylene activates in AZs to orchestrate the separation of floral organs. AZ promoter activity of *IDA* was indeed found to depend on ethylene signaling (Butenko et al., 2006). Wounding was also shown to induce early activation of *IDA* promoter in AZs (Butenko et al., 2006). In cauline AZs, drought stress induces *IDA* transcription (Patharkar and Walker, 2016). In the lateral root emergence zone of seedlings, *IDA* promoter activity is enhanced in response to the Microbe-Associated Molecular Patterns (MAMPs) flagellin (flg22), chitin, as well as to salt and mannitol (Lalun et al., 2023). These instances suggest that a sizeable set of environmental factors known to influence abscission are integrated in the transcriptional regulation of *IDA*.

In this study, we investigate the genetic and molecular determinants of *IDA* transcriptional regulation. An *in silico* dissection of the *IDA* promoter and a screen against an Arabidopsis TF collection highlight the diversity of cues and effectors that can modulate *IDA* expression. We demonstrate that DNA binding sites of ethylene response factors (ERFs) and WRKY TFs are required for full transcriptional competence of *IDA* and to mediate its MAMP-dependent transcriptional upregulation, respectively. We also show that WRKY57 can modulate floral organ abscission in an IDA/IDL and HAE/HSL2-dependent manner.

## Materials and methods

### Plant material, growth conditions and treatments

All Arabidopsis lines were in the Columbia (Col-0) genetic background, except for the *ida-1* mutant (C24; (Butenko et al., 2003)). The previously published mutant and transgenic lines used in this study were: *ida-2* (Cho et al., 2008), *idaCR* (Alling and Galindo-Trigo, 2023), *idl1CR* (Shi et al., 2018), *p35S::IDA* (Stenvik et al., 2006), *hae hsl2* (Stenvik et al., 2008), *wrky57* (Jiang et al., 2012), *wrky60-1* (Xu et al., 2006) and *wrky48* (Jiang et al., 2012). Genotyping primers are listed in the Supplemental Table S1.

Arabidopsis plants were routinely vapor-sterilized with chlorine gas, sown on Murashige & Skoog (MS) medium plates with 0.7% sucrose, stratified for three days and germinated in growth chambers for a week before transfer to regular sowing soil. Subsequently, plants grew in climate rooms until seed setting and senescence. Environmental growth conditions in the growth chamber and climate rooms were similar: photoperiod of 16h day/8h night, light intensity of 130-150 μmol·m^-2^·s^-1^, temperature of 22°C, and 60% humidity. Transgenic plants were selected in plates supplemented with Basta or hygromycin-B as required. Microscopy of roots was carried out with vertically grown seedlings in 0.5xMS plates with 0.7% sucrose.

In the case of the flg22-treated mature rosette leaves, seeds were directly germinated on peat pellets (Jiffy 7) with short day photoperiod (10h day/14h night). Two plants per pellet were allowed to progress past seedling stage. At week 6 and prior to bolting, the most expanded rosette leaf of each plant was syringe infiltrated with mock solution (water) or flg22 solution (500 nM flg22 in water). Twenty hours later, leaves were detached and individually processed to detect GUS activity. Leaves were assigned to different qualitative categories with values ranging from 0 (undetectable GUS staining) to 3 (strong GUS signal in vasculature and neighboring leaf tissues). To quantify the effect of flg22 on root meristems, seedlings were germinated in liquid MS as per (Luna et al., 2011). Liquid medium was refreshed after one week of growth, and flg22 treatments applied one day later. Mock treatments (water) or flg22 treatments (1 µM flg22 in water) were applied in the evening of the 8^th^ day, and seedlings processed for microscopy the following day after approximately 20h of treatment. Quantification of *IDA* induction in root meristems was conducted by producing maximum intensity projections of the H2B-TdTomato channel images, and counting the total number of fluorescent nuclei in the meristematic region using the StarDist plugin in ImageJ with predetermined settings (Schmidt et al., 2018). To assay the responsiveness of AZs to flg22, plants were grown to maturity in standard conditions. Developing siliques in positions 6-7 that had already shed all floral organs were selected and separated from the plant by the pedicel. The siliques were immediately submerged in the mock (water with 0.02% Silwet L-77) or flg22 treatments (10 µM flg22, 0.02% Silwet L-77 in water) for 15 minutes. Subsequently, the siliques were transferred to the overnight incubation solutions for mock (water) or flg22 treatments (10 nM flg22 in water). The initial short treatment with detergent allows for the explants to become less hydrophobic and an even elicitation. Incubation overnight in the solutions without detergent helps avoid toxicity. Cauline leaf elicitation with flg22 was conducted on five-week-old plants grown under standard conditions. The first two cauline leaves of each plant were syringe-infiltrated with mock solution (water) or flg22 solution (1 µM flg22 in water). The entire surface of the leaf was infiltrated, including the boundary between leaf and stem. The treatment was allowed to proceed for 20h. The cauline leaf-stem boundary was manually dissected with a razor blade and imaged with confocal microscopy immediately after.

*Nicotiana benthamiana* plants were grown in growth chambers with a long day photoperiod, diurnal temperature of 22°C and night temperature of 19°C, 150-180 μmol·m^-2^·s^-1^ of light, and 60% relative humidity.

### Generation of genetic constructs and transgenic lines

Arabidopsis plants were transformed with Agrobacterium C58 following the floral dip method (Clough and Bent, 1998).

Most genetic constructs used in this study were generated with Invitrogen Gateway recombination cloning. Promoter sequences were cloned into the cloning vector pENTR5’ (TOPO-TA cloning; Invitrogen). The *IDA* promoter was cloned from C24 gDNA as the 1417bp between the *IDA* translation initiation site (TIS) and the upstream gene *AT1G68780*. *IDA* promoter sequences in C24 and Col-0 accessions are identical, with the exception of the length of a dinucleotide repeat located 144bp from the TIS that is extended by 12bp (CACACACACAAG) in the C24 genome. In all other cloning instances, the Col-0 accession was used as template. The *ERF*(-) and *WRKY*(-) versions of the *IDA* promoter were generated by site-directed mutagenesis (Zheng et al., 2004). Five consecutive rounds of site-directed mutagenesis were necessary to generate the *WRKY*(-) *IDA* promoter. The *IDL1* promoter comprises 1557bp between its TSS and the gene upstream AT3G25660. The IDL2 promoter comprises 2084bp upstream of its TIS. The *IDL3* promoter comprises 1974bp upstream of its TIS. *WRKY57* promoter covers 2071bp upstream of its TIS. *WRKY60* promoter comprises the 1512bp between its TIS and the upstream gene *AT2G24990*. The *HAESA* promoter contains 1729bp upstream of the 37^th^ bp from the TIS. Coding sequences to be expressed *in planta* were cloned into pDONR221 or pDONR-Zeo in BP Clonase II reactions (Invitrogen). Intronless coding sequences from *IDA*, *WRKY57* and *WRKY60* were PCR amplified from floral tissue cDNA – see the RT-qPCR section for methods regarding RNA extraction and cDNA synthesis. The WRKY57srdx entry vector was obtained by amplifying *WRKY57* coding sequence from floral cDNA using a modified reverse primer containing the coding sequence of the repressor motif SRDX (CTCGATCTGGATCTAGAACTCCGTTTGGGTTTCGCT) in frame with the C-terminus of *WRKY57*. These and entry vectors were recombined into destination vectors from the Nakagawa lab (Nakagawa et al., 2007; Nakagawa et al., 2008; Tanaka et al., 2013) by means of LR recombination reactions using either LR II Clonase or LR II Clonase Plus (Invitrogen). The destination vector used to generate *GFP-GUS* promoter reporter lines was R4L1pGWB632. Luciferase promoter reporters were generated with the destination vector R4L1pGWB635. GFP-tagged translational fusion reporters were cloned into R4pGWB504. Non C-terminally tagged *promoter*:*cds* constructs were cloned into R4pGWB501. Effector constructs to overexpress TFs or negative controls in luciferase assays were cloned into pGWB518.

To generate nuclear fluorescent promoter reporters *H2B-TdTomato* and *Venus-H2B*, a pDONR221 entry clone containing the coding sequence of the fusion proteins H2B-TdTomato or Venus-H2B were first generated. The *H2B-TdTomato* sequence was PCR amplified from pAH21-H2B-TdTomato, while the *Venus-H2B* sequence was amplified from pAB146 (kindly provided by Simon Rüdiger). These entry clones were subsequently recombined with the corresponding promoter and/or destination vectors.

Col-0 wild-type plants were gene edited to produce the *idl2CR* and *idl3CR* lines with the plasmid system by Fauser et al. comprising pDe-Cas9 and pEn-Chimera vectors (Fauser et al., 2014). Guide RNA protospacers targeting *IDL2* in its 69^th^ codon (BmgBI restriction site), and *IDL3* in its 37^th^ codon (XmnI restriction site) were designed. Transformants were screened by CAPS and mutant alleles confirmed by Sanger sequencing. A +1bp insertion causes a frameshift immediately prior to the IDL2 peptide in *idl2CR* plants, whereas in *idl3CR* plants a −1bp deletion causes a frameshift in the variable region of the IDL3 protein. The Cas9 T-DNA cassette was segregated away from the plant genetic background and homozygous plants for the *idl2CR* or *idl3CR* mutations were confirmed by CAPS and sequencing.

Constructs used in the yeast one-hybrid (Y1H) assay were generated by recombining in Gateway LR reactions the *IDA* promoter entry vector with the two yeast reporter plasmids - pMW2 and pMW3 (Deplancke et al., 2006) - containing respectively HIS3 or LacZ reporter genes.

Primers used to generate these constructs can be found in the Supplemental Table S1.

### Detection and visualization of expression reporters

Promoter activity in beta-glucuronidase (GUS; (Jefferson et al., 1987)) reporter lines was detected as follows: explants/seedlings were incubated in ice-cold 90% acetone for 20 minutes, washed for 20 minutes in staining buffer without X-Gluc (50 mM NaPO_4_ pH 7.4, 2 mM potassium ferro-cyanide, 2 mM potassium ferri-cyanide, 0.1% Triton X-100), and stained in staining buffer with 2 mM X-Gluc at 37°C for 3h or overnight. The chlorophyll was then cleared from the tissues with washes in 75% ethanol for 1-3 days and imaged on a stereomicroscope or widefield microscope.

Fluorescent reporter detection was conducted on live tissue. Explants were mounted in water and immediately imaged in a Zeiss LSM880 confocal microscope. To visualize GFP-tagged translational fusions as well as the GFP-GUS dual reporter or Venus-H2B, fluorophores were excited with a laser wavelength of 488nm. Nuclear transcriptional reporters H2B-TdTomato were excited with a wavelength of 561nm. To colocalize nuclear reporters (GFP or H2B-TdTomato) with 4′,6-diamidino-2-phenylindole (DAPI), live explants were incubated in DAPI staining solution (0.2 mg/L in water) for 30 min and subsequently imaged with an excitation wavelength of 405nm. Chlorophyll autofluorescence was excited with a laser wavelength of 633 nm to allow easier identification of AZ or cauline leaf regions.

### AZ and cell size estimations

AZs of six-week-old plants grown in standard conditions were imaged in a stereomicroscope. The transversal area occupied by the receptacle was manually selected in ImageJ (Schneider et al., 2012), and then measured. To estimate the cell size in AZs, the AZs were stained with propidium iodide (10 mg/L in water) and imaged in a Zeiss LSM880 confocal microscope with laser excitation at 561nm. Concomitantly, the GFP-tagged proteins were imaged with excitation wavelength of 488nm. For each AZ, the longest possible diameter of five consecutive cells from the sepal AZ were measured manually with ImageJ and then averaged to produce the mean estimated cell size of an AZ. Transversal AZ area was measured in position 12 siliques. Cell size was measured in position 8. These positions displayed the greatest contrast in size between wild-type plants and genotypes in which AZs enlarged, while technically allowing the imaging to be conducted reliably.

### Gene expression estimations by RT-qPCR

Floral receptacles were manually dissected from at least 25 flowers (positions five/six) collected from three to four plants to yield each biological replicate. Excised receptacles were immediately transferred to tubes pre-chilled in liquid nitrogen, and flash-frozen in liquid nitrogen. RNA was extracted using either Spectrum Plant Total RNA Kit (Sigma) or RNeasy Plant Mini Kit (Qiagen) following the manufacturer instructions and including an on-column DNase I digestion step (Sigma). First strand cDNA was synthesized with Superscript III or Superscript IV Reverse Transcriptase (Invitrogen), RNA digested with RNase H, and selected loci amplified and quantified with FastStart Essential DNA Green Master (Roche) in a LightCycler96 instrument (Roche). Gene expression was estimated with the 2^-ΔCt^ method using *ACTIN2* as a reference gene. Experiments comprised three biological replicas per genotype and two technical replicas per RT-qPCR reaction. Primers used in RT-qPCR reactions can be found in the Supplemental Table S1.

### Luciferase promoter transactivation assays

Transactivation assays in transiently transformed *N. benthamiana* plants were carried out as per (Lasierra and Prat, 2018). Briefly, fully expanded leaves from four-week-old plants were syringe-infiltrated with Agrobacterium solutions carrying the reporter and effector plasmids at 0.02 OD_600_ each. Measurements were conducted three days after infiltration on a 96-well OptiPlate (PerkinElmer) by floating 4 mm diameter leaf discs (abaxial side up) on 200 µL of luciferin solution (1xMS salts, 0.5% MES, pH 5.8, 12 µM D-luciferin). Light emission was measured in a Wallac 1420 VICTOR2 microplate reader (PerkinElmer) recording the light emitted for 10 seconds, with 10 minutes delay between each repeat. The third plate repeat after approximately 35 minutes typically recorded the strongest signal and was used to plot and analyze differences between constructs. Twelve leaf discs per construct combination coming from two different plants per construct combination were assayed per experiment.

Assays with the *IDA* promoter luciferase constructs were first attempted in *N. benthamiana* leaves but strong autoactivation of the *IDA* promoter in this system impeded reliable quantifications. An Arabidopsis mesophyll protoplast transient transfection system was used instead. We used fully expanded rosette leaves from pre-bolting, four-week-old Col-0 plants grown in our standard conditions to extract the protoplasts following the tape-sandwich method (Wu et al., 2009; Hansen and van Ooijen, 2016). Transfections were carried out with 50 µL of protoplasts at 400000 protoplasts/mL and 6 µg of plasmid DNA, purified with PureLink HiPure Plasmid Midiprep Kit (Thermofisher). After transfections, protoplasts were allowed to recover in W5 solution overnight. The protoplasts were gently pelleted and resuspended in 100 µL of W5 solution with 1 mM D-luciferin and immediately transferred to 96-well plates. Light emission was recorded in for five seconds, with eight minutes delay between each repeat. Four independent protoplast transfections per construct combination were analyzed per experiment.

### *In silico* analyses of promoter sequences

The promoter sequence of *IDA* was scouted for presence of TFBSs using Binding Site Prediction tool from the Gao lab’s PlantRegMap site (http://plantregmap.gao-lab.org/; Supplemental Table S2; (Tian et al., 2019)). Gene ontology term enrichment analysis was carried out using the GO Term Enrichment tool in PlantRegMap. The input was the list of TFs identified as *IDA* promoter interactors in either HIS3 or LacZ assays, and the reference set of genes was the total list of TFs from Arabidopsis thaliana – downloaded from the PlantRegMap site. This tool calculates statistically significant enrichment with topGO and Fisher’s exact tests, with threshold p-value ≤ 0.01. Additional TFBS searches were conducted with the PlantPAN 3.0 web tool (http://plantpan.itps.ncku.edu.tw/plantpan4; (Chow et al., 2019)).

### Yeast one hybrid screen

The *IDA* promoter was recombined in the pMW2 and pMW3 plasmids, respectively, to drive expression of the HIS3 and LACZ reporters (Gaudinier et al., 2017). These reporter plasmids were transformed into the yeast YM4271 strain and the yeast colonies were screened for autoactivation and the construct presence via PCR genotyping. The Enhanced Yeast One-Hybrid screening of the *IDA* promoter against a complete collection of 2000 Arabidopsis transcription factors was done as described previously (Gaudinier et al., 2011; Pruneda-Paz et al., 2014; Gaudinier et al., 2017). The positive interactions were recorded for LacZ and HIS3 activity. The Yeast One-Hybrid screening was carried out by the Yeast One Hybrid Services Core at the UC Davis Genome Center, at the University of California, Davis (https://genomecenter.ucdavis.edu/yeast-one-hybrid-services). Y1H screening results are listed in the Supplemental Table S3.

### Floral organ retention quantification

To phenotype and quantify abscission, plants were grown in individual pots and plants from different genotypes shuffled in their positions across the growth tables to minimize positional effects. Plants were grown undisturbed and untouched to minimize uneven shedding of floral organs prior to phenotyping. Phenotyping was generally conducted at week 6 when inflorescences had produced between 20-25 flowers post anthesis. Floral organ abscission was quantified by counting the floral organs attached to the flowers in the main inflorescence as per (Alling and Galindo-Trigo, 2023). The main inflorescence stem was shaken four times and the number of floral organs that remained attached to each floral position (P1-P20) was visually inspected.

## Results

### Dissection of promoter cis elements suggests the *IDA* gene is subject to intricate transcriptional regulation

One of the main genomic features that dictate the expression profile of a gene is the presence of cis regulatory sequences to which TFs specifically bind (TF binding sites (TFBSs)). We used the web based PlantTFDB 4.0 database to determine the presence of conserved TFBSs in the promoter of *IDA* to investigate the determinants of its transcriptional regulation (Jin et al., 2017). We detected TFBSs for all the major TF families in plants, including ethylene-response factor (ERF), and WRKY, among others (Supplemental Figure S1 and Supplemental Table S2). The majority of the TF families detected presented one or more TFBSs within 500 nucleotides of the translation initiation site (TIS) of *IDA*, the portion of promoters shown to withstand the most stringent evolutionary constraints in a panel of Arabidopsis accessions (Korkuc et al., 2014). This suggests the transcription of *IDA* may be effectively controlled by an extensive array of TFs, allowing for its spatially and temporally-restricted, yet highly environmentally-responsive expression pattern (Butenko et al., 2003; Butenko et al., 2006; Vie et al., 2015; Patharkar and Walker, 2016; Lalun et al., 2023).

Given the extensive list of potentially important TFBSs (Supplemental Table S2), we decided to conduct proof-of-concept experiments on ERF and WRKY binding sites to functionally demonstrate the physiological relevance of TFBSs for *IDA* expression. Members of the ERF and WRKY TF regulate physiological processes adjacent to *IDA* signaling and abscission such as ethylene signaling, drought responses, MAMP-induced responses, or senescence (Lorenzo et al., 2003; Zhou et al., 2011; Jiang et al., 2012; Chang et al., 2013; Cheng et al., 2013; Lyons et al., 2013; He et al., 2016; Jiang and Yu, 2016; Lal et al., 2018). We used site-directed mutagenesis to disrupt selected TFBSs for ERFs and WRKYs in the *IDA* promoter, and observed the effect of the mutations on the expression of the promoter with a GUS-GFP dual reporter in stably transformed Arabidopsis plants (Tanaka et al., 2013). Disruption of a single ERF TFBS predicted to convey signals of up to 66 ERF TFs was sufficient to reduce the activity of the *IDA* promoter to barely detectable levels in AZs and floral organs (*ERF*(-); Figure 1A, B and Supplemental Figure S2). The *ERF*(-) *IDA* promoter was also inactive during lateral root emergence (Supplemental Figure S3A). We then tested its capacity to genetically complement the abscission phenotype of *ida* knock-outs (Butenko et al., 2003; Cho et al., 2008). The *ERF*(-) promoter was unable to revert the abscission phenotype of *ida*, indicating that its weak activity in nectaries does not induce floral organ separation (Figure 1C, Supplemental Figure S4). These results indicate the ERF-mediated signaling is crucial for *IDA* expression and required for abscission to take place. On the other hand, disrupting the three WRKY TFBSs predicted with our initial search did not yield noticeable changes to the promoter activity (Supplemental Figure S1). Additional WRKY TFBSs were found in the *IDA* promoter by alternative bioinformatic tools and a quantifiable decrease in the promoter activity in AZs was observed when five binding sites were mutated (*WRKY*(-); Figure 1A, B and Supplemental Figure S2 and S5; (Chow et al., 2019)). Despite its decreased activity in AZ cells, the *WRKY*(-) *IDA* promoter was still active in floral organs and during lateral root emergence, suggesting that WRKY TFs have an important but not essential role on the developmentally-induced expression of *IDA* (Supplemental Figure 3B).

**Figure 1.**
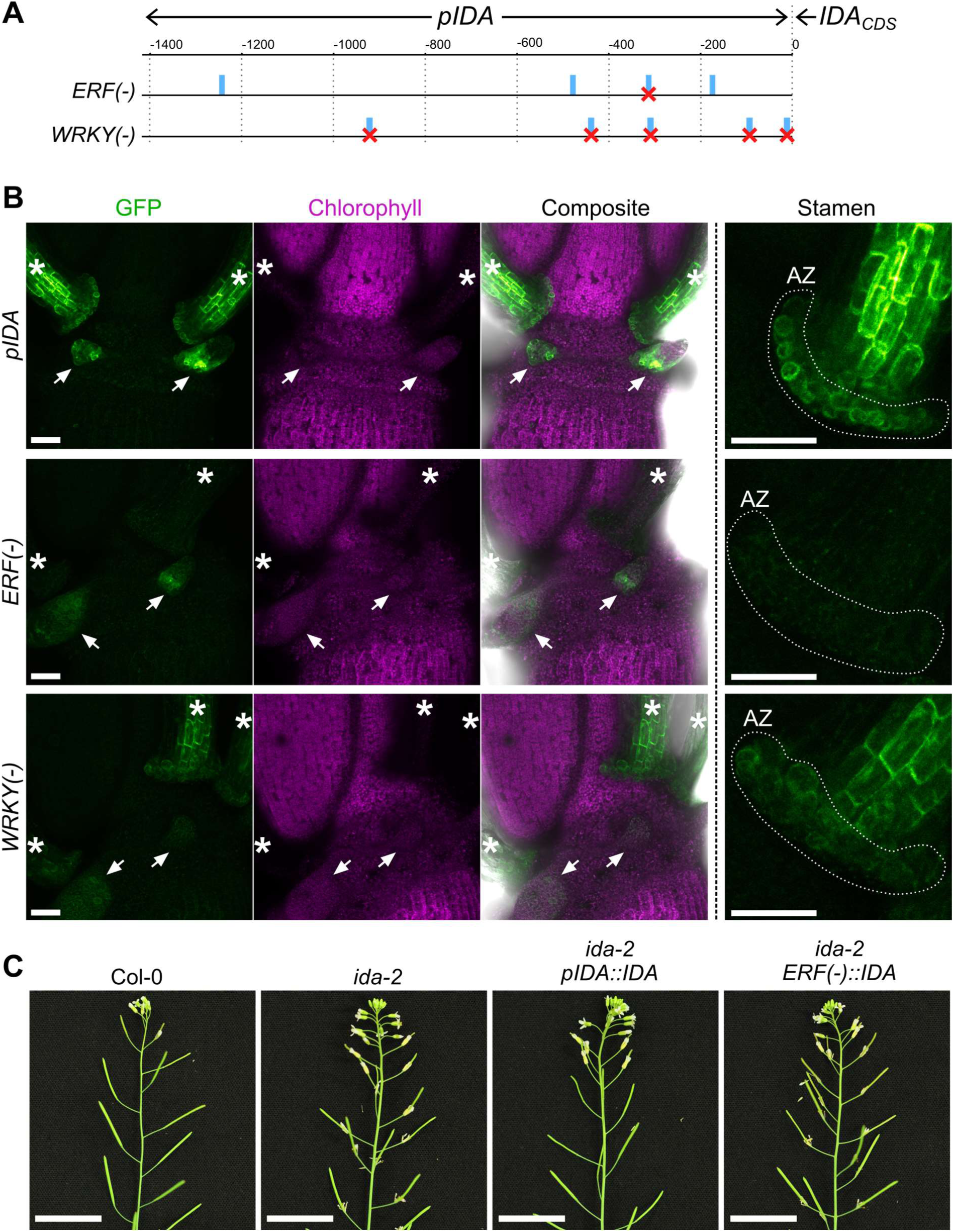
The *IDA* promoter activity is subject to distinct transcriptional regulation by ERF and WRKY transcription factors. **A**, Schematic diagram of the *IDA* promoter sequence and the two mutant versions: *ERF*(-) and *WRKY*(-). Numbers in the upper scale represent the distances in number of nucleotides from the translation initiation site of the *IDA* gene. Blue rectangles indicate the predicted TFBSs for ERF or WRKY TFs along the *IDA* promoter. Crosses mark the specific TFBSs that were disrupted by site-directed mutagenesis in either version of the *IDA* promoter – One ERF binding site, and five WRKY binding sites, respectively. **B**, Representative confocal micrographs of plants expressing GFP-GUS. Each row corresponds to a promoter reporter line, indicated on the left. The GFP is shown in green, chlorophyll autofluorescence is shown in magenta. The rightmost column shows detail images of the stamen proximal region and AZ. White dotted lines highlight the AZ region from which quantitative measurements in Supplemental Figure S2B were taken. White arrows point towards nectaries. White stars highlight stamens. Scale bars at both magnifications represent 50 µm. **C**, Floral organ abscission phenotypes of *ida-2* mutants genetically complemented with the *IDA* gene under its wild-type promoter (*pIDA*) or the *ERF*(-) mutant version. Floral organs remain attached to most developing siliques in the *ERF*(-) lines, indicating lack of complementation of the *ida-2* abscission defect. Twelve out of 14 *pIDA* lines fully reverted the *ida-2* phenotype. Zero out of 20 *ERF*(-) lines fully reverted the *ida-2* phenotype. See also Supplemental Figure S4. Scale bars are 2 cm.

### WRKY binding sites are necessary to activate the *IDA* promoter in response to the immunity elicitor flg22

When plants are compromised by infections, organs that would otherwise be retained are abscised to protect the plant (Lahey et al., 2004; Scalschi et al., 2014; Patharkar et al., 2017). It is well established that bacterial and fungal elicitors induce *IDA* expression, however the molecular mechanism driving this transcriptional upregulation is not known (Vie et al., 2015; Lalun et al., 2023). Several WRKY TFs are elicited by MAMPs like flg22, are capable of inducing transcription of defense-related genes, and their overexpression enhances disease resistance (Asai et al., 2002; Navarro et al., 2004; Zipfel et al., 2004; Birkenbihl et al., 2016). We thus hypothesized WRKY TFBSs in the *IDA* promoter could influence its responsiveness to pathogenic cues. Indeed, *WRKY*(-) reporter lines were significantly less responsive than lines with wild-type *IDA* promoter to treatments with flg22 in multiple tissues (Figure 2). Wild-type *IDA* promoter was strongly activated by flg22 in AZ cells, whereas its *WRKY*(-) counterpart only showed a weak induction (Figure 2A and B). In seedlings, the flg22 treatment also induced the *IDA* promoter in the meristematic zone of the main root, while the *WRKY*(-) IDA promoter failed to do so (Figure 2C and D). The same result was observed when fully expanded rosette leaves were infiltrated with flg22, as the observed induction of *IDA* promoter along the leaf midrib was reduced in the *WRKY*(-) reporter lines (Figure 2E and F). The receptors of IDA, HAE and HSL2, are involved in pathogen-induced cauline leaf abscission in Arabidopsis (Patharkar et al., 2017). We therefore looked for upregulation of the *IDA* promoter in cauline abscission zones after infiltrating cauline leaves with flg22. We did not detect *IDA* promoter activity in this tissue under our conditions (Supplemental Figure S6). Our results support a role for WRKY TFs in mediating *IDA* expression upon biotic stress. We propose the *IDA* promoter to be a direct target of immunity-activated WRKYs (Asai et al., 2002; Lal et al., 2018).

**Figure 2.**
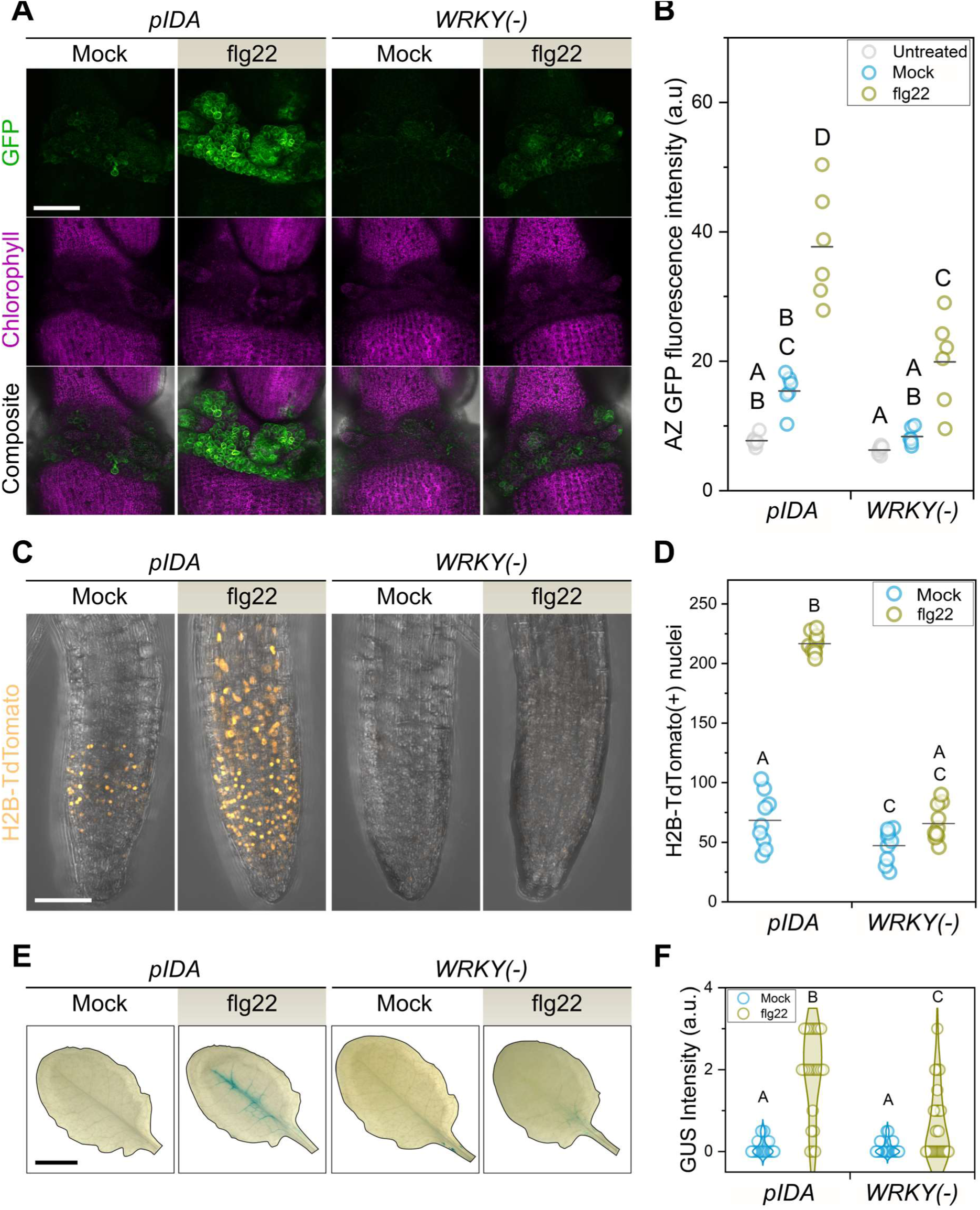
*IDA* responsiveness to flg22 depends on the presence of WRKY promoter binding sites. **A**, Residuum abscission zone cells induce *IDA* in response to flg22 in a WRKY TFBS-dependent manner. Reporter lines express GFP-GUS under the control of the promoters indicated on top. Scale bar represents 120 µm. Micrographs shown are maximum intensity projections of the GFP, chlorophyll autofluorescence or composite channels. Images are representative of an experiment with six plants per line, and one flower in positions six or seven per plant per treatment. The experiment was repeated twice using different transgenic lines for each reporter construct with similar results. **B**, Quantification of the GFP fluorescence intensity in the entire abscission zone area of the maximum intensity projections shown in A. Each datapoint corresponds to the average GFP intensity in the abscission zone region of one flower. **C**, The root meristematic region responds to flg22 treatment by upregulating *IDA* in a WRKY TFBS-dependent manner. Reporter lines express H2B-TdTomato under the control of the promoters indicated on top. Images are maximum intensity projections of composite confocal micrographs. Scale bar represents 30 µm. Images are representative of an experiment with ten seedlings per line and treatment. The experiment was repeated three times with three independent lines per reporter construct and obtained similar results. **D**, Quantification of nuclei with H2B-TdTomato expression from maximum intensity projections of the root meristematic region. Each datapoint corresponds to the nuclei quantified in each individual seedling assayed. **E**, Representative images of the histochemical detection of GUS activity in mature rosette leaves in plants expressing GFP-GUS under the promoters displayed on top. Scale bar is 0.5 cm. **F**, Quantification of the beta glucuronidase (GUS) activity detected in mature leaves like the ones shown in E. Data shown here contains the dataset from three independent transgenic lines per reporter construct, totaling between 24-32 plants per construct and treatment combination. Leaf GUS staining intensity was qualitatively assessed and assigned to categories (see Materials and methods). Letters in panels B, D and F represent categories of statistically significant differences in two-way ANOVA and post hoc pairwise comparisons with Bonferroni tests (p < 0.05).

### WRKY57 is a potential regulator of *IDA* transcription

Next, we aimed to identify and functionally characterize WRKYs that regulate *IDA* expression during floral organ abscission. We screened a collection of 2000 TFs of Arabidopsis in a yeast one-hybrid (Y1H) assay to identify those that bound the *IDA* promoter. This screening detected interaction between the *IDA* promoter and 211 TFs, the majority of which belonged to the main TF types predicted with the *in silico* analysis (Figure 3A and Supplemental Figure S1). A gene ontology enrichment analysis of the 211 TFs that bind the *IDA* promoter revealed enriched categories expected of abscission-related genes: floral whorl morphogenesis and carpel formation, sugar and carbohydrate-mediated signaling, regulation of ethylene responses, among others (Figure 3B). Twelve WRKYs were detected in the Y1H screening and most of them were induced in abscission zones as abscission progresses (Supplemental Figure S7A). Out of the 12, WRKY57 and WRKY60 were selected as our primary candidates to regulate *IDA* because i) their own upregulation preceded the induction of *IDA*, and ii) *WRKY57* and *WRKY60* had been highlighted in the list of most significantly regulated genes of the AZ transcriptome (Cai and Lashbrook, 2008). Besides, WRKY57 is involved in balancing jasmonic acid and auxin signaling in leaves during senescence, is a negative regulator of biotic stress resistance and its overexpression confers drought tolerance – processes previously linked to IDA signaling and abscission (Jiang et al., 2012; Jiang et al., 2014; Jiang and Yu, 2016; Serrano-Bueno et al., 2022). Similarly, WRKY60 is involved in immunity, and ABA signaling in osmotic and salt stress (Xu et al., 2006; Chen et al., 2010; Liu et al., 2012).

**Figure 3.**
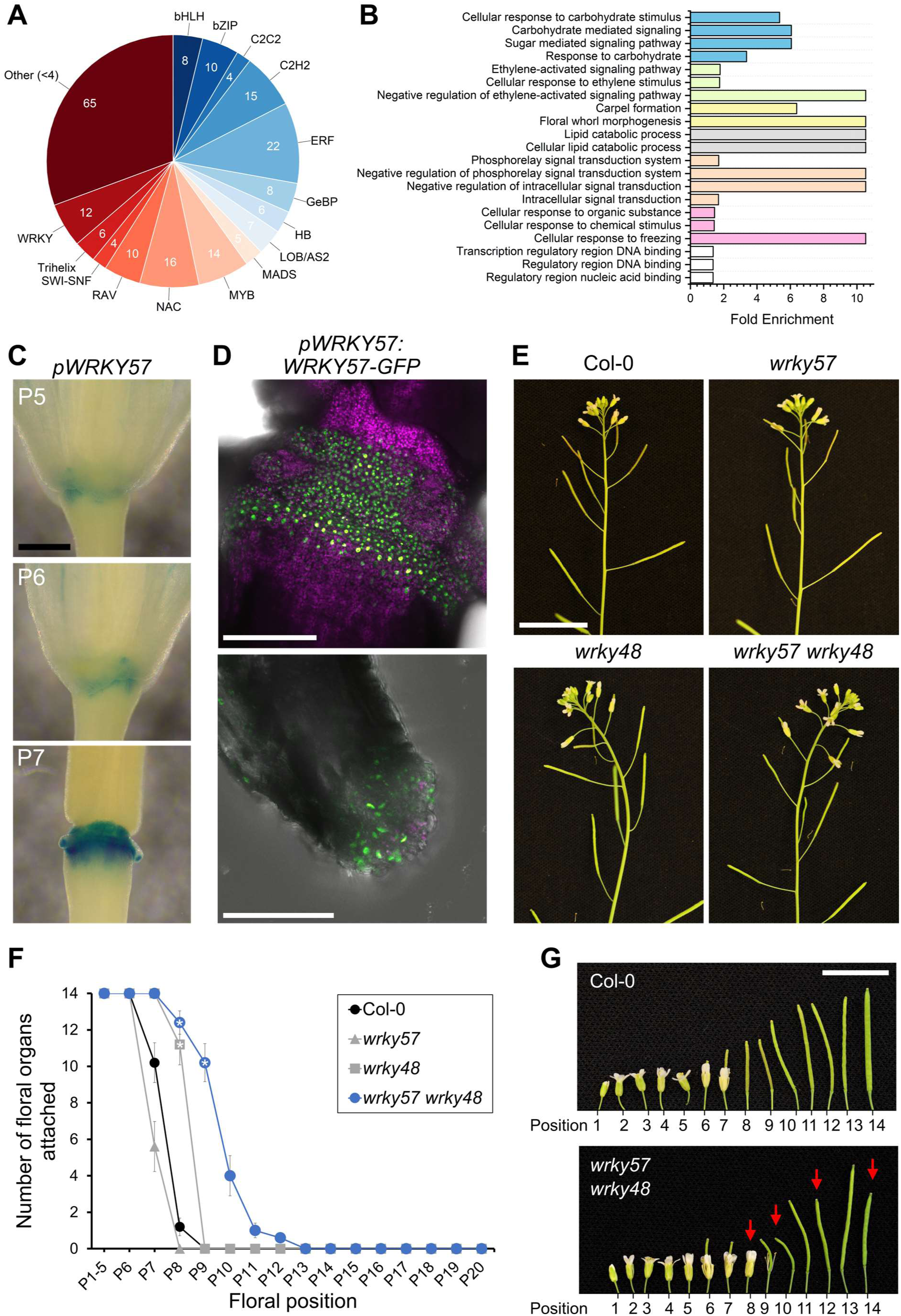
A screening against a collection of Arabidopsis TFs identified WRKY57 as a putative regulator of abscission. **A**, Summary of TFs identified in the Y1H screen of the *IDA* promoter. TFs are represented by family. TF families with less than 4 members identified in the screen were grouped in the “Other” category to ease visualization. See also the Supplemental Table S3. **B**, Gene ontology term enrichment analysis of the set of 211 TFs identified in the Y1H screen. Gene ontology terms associated with the 211 TFs were compared against the reference, in this case the entire list of TFs in Arabidopsis thaliana. All terms listed here are statistically enriched at p < 0.01. Different colors represent related terms (blue, carbohydrates; green, ethylene; yellow, floral development; grey, lipids; orange, signaling; pink, other responses; white, DNA binding). **C**, Histochemical staining of GUS activity in plants expressing GFP-GUS under *WRKY57* promoter. Scale bar is 0.4 mm. P5, P6, P7 are floral positions five, six and seven along the main inflorescence stem. **D**, Maximum intensity projections of confocal micrographs of the AZs of plants expressing *pWRKY57:WRKY57-GFP*. On the top panel, green nuclear signals from WRKY57-GFP are seen in the residuum cells. Scale bar is 150 µm. Below, WRKY57-GFP accumulates in the secession cells of a petal. Scale bar is 50 µm. Chlorophyll autofluorescence is shown in magenta in both images. **E**, Inflorescences of T-DNA lines *wrky57*, *wrky48*, and double mutant *wrky57 wrky48*. Scale bar is 2 cm. **F**, Quantitative phenotyping of floral organ abscission in the mutant lines listed in the graph legend. Markers (circles, squares, triangles) represent the mean, and whiskers the SEM. Data corresponds to the number of floral organs fully attached to the flower at each position from five plants per genotype. One way ANOVA and post hoc Bonferroni test were used to compare the four genotypes at each position. Positions with a statistically significant difference in the mean compared to wild-type plants are highlighted with a white star (p < 0.05). This assay was repeated twice with similar outcomes. **G**, Irregular self-pollination in the *wrky57 wrky48* double mutant, highlighted by red arrows. Scale bar is 1 cm.

To verify the transcriptomic data *in planta*, transcriptional and translational fusion reporter lines were generated. Unexpectedly, the reporter lines of *WRKY60* indicated that both its promoter activity and its protein accumulation take place outside of the AZs, where WRKY60-GFP accumulates in the nucleus (Supplemental Figure S7B, C and S8A). A *wrky60* T-DNA insertion line was phenotyped at the flowering stage and no abscission defect was observed (Supplemental Figure S7D). Based on this set of results, we discarded WRKY60 as a putative regulator of *IDA* in abscission. The promoter of *WRKY57* was active in AZs prior to abscission, and WRKY57-GFP accumulated in the nuclei of AZ cells, supporting a putative role for WRKY57 in regulating *IDA* and abscission (Figure 3C and D, and Supplemental Figure S8B). A single mutant *wrky57* abscised as wild-type however, WRKYs are considered to be largely redundant with other members of their family (Figure 3E; (Du et al., 2023)). Functional redundancy is most common between proteins with the highest homology and thus we generated a double knock-out mutant between *WRKY57* and *WRKY48,* WRKY57’s closest homolog in BLAST searches against the Arabidopsis proteome (Altschul et al., 1990; Jiang et al., 2012). A delay in abscission was observed in *wrky57 wrky48* double mutants compared to wild-type, suggestive of a weak abscission defect (Figure 3E and F). It must be noted that *wrky57 wrky48* mutants fail to fully fertilize some of their siliques (Figure 3G), and so caution should be taken when concluding the cause of the abscission delay observed in this double mutant. Collectively, these findings portrayed WRKY57 as a potential regulator of *IDA* and abscission in redundancy with other WRKYs. This prompted us to functionally characterize WRKY57 activity in abscission zones.

### Overexpression of *WRKY57* or a *WRKY57* repressor respectively activates or represses floral organ abscission

TFs can positively or negatively regulate the transcription of genes. Typically, TFs bind TFBSs in the promoter region of genes, and recruit additional effectors to mediate activation or repression of transcription. WRKY57 was shown to induce or repress jasmonic acid-induced senescence in Arabidopsis leaves by competitively interacting with repressors from the JASMONATE ZIM-DOMAIN (JAZ), or AUX-IAA families, respectively (Jiang et al., 2014). This functional duality indicates that the activity of WRKY57 is likely dependent on the tissue, developmental stage and environmental cues, as does the abundance of interactors and hormones that influence WRKY57. To shed light on the role of WRKY57 during abscission we locally overexpressed *WRKY57* using the *HAE* promoter – highly active during abscission in both residuum and secession cells (Jinn et al., 2000; Lee et al., 2018). Multiple transgenic lines overexpressing WRKY57-GFP in AZs displayed enlargement of the receptacle after abscission, correlating with the transgene level of expression (Figure 4A, B and Supplemental Figure S9A, B). The AZ enlargement was caused by excessive cell expansion, reminiscent of that observed in plants constitutively overexpressing *IDA* (*p35S*::*IDA*; (Stenvik et al., 2006)), and those expressing *IDA* under the *HAE* promoter (*pHAE*::*IDA*; Figure 4C and Supplemental Figure S9C). The excessive cell expansion is likely due to overactive cell separation, as loose cells detach from the AZs of the WRKY57-GFP overexpressing lines when mounting samples for microscopy (Figure 4D).

**Figure 4.**
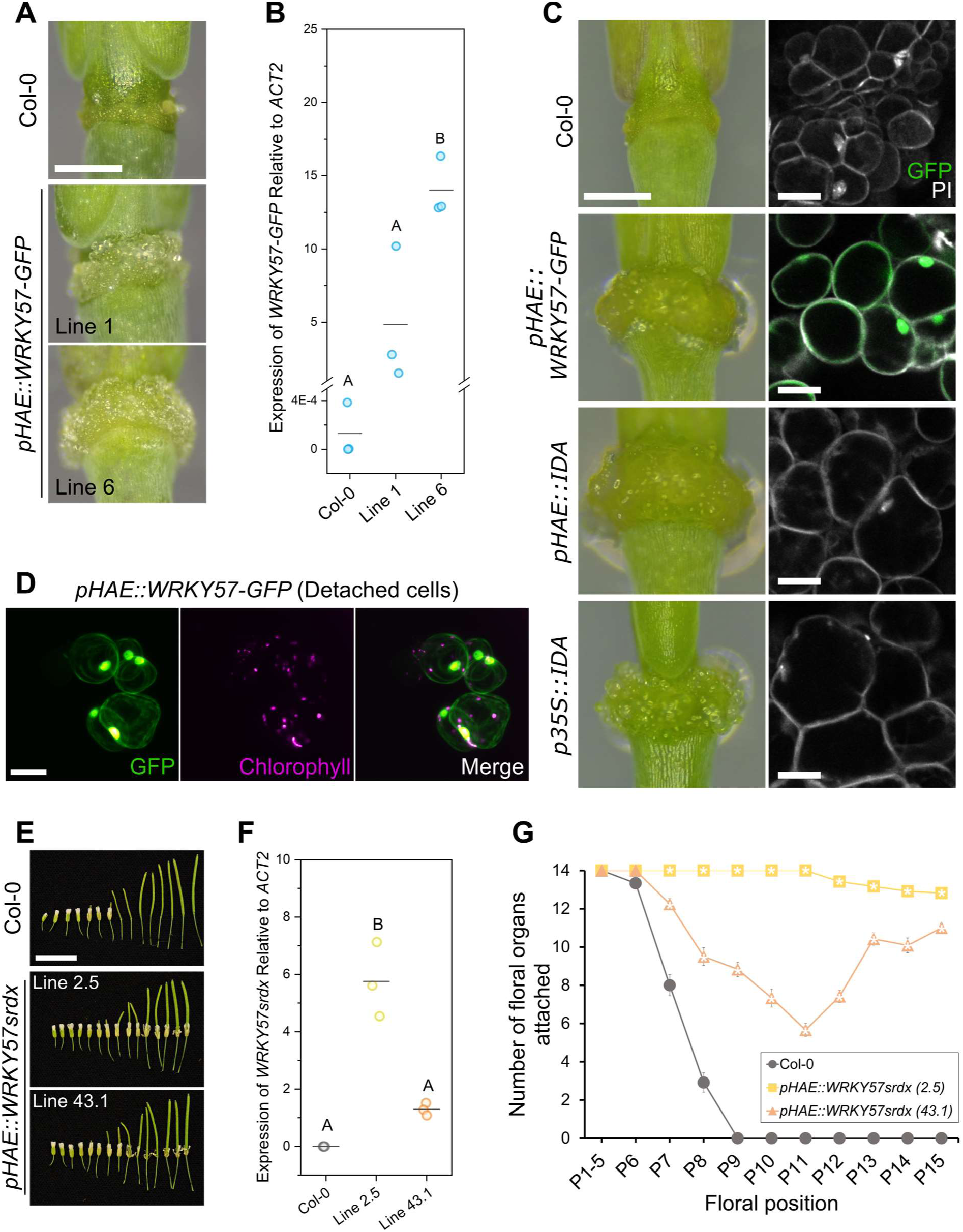
Modulation of floral organ abscission by local overexpression of *WRKY57*. **A**, Floral receptacles of transgenic lines in Col-0 background overexpressing WRKY57-GFP in floral position 12. Scale bar is 0.4 mm. **B**, Quantification of transgene expression levels in the transgenic lines shown in A. Datapoints are one biological replica (averaged from two technical replicas). Three biological replicas were analyzed per line. Letters denote statistically different groups at p < 0.05 in one way ANOVA and Bonferroni post hoc tests. *ACT2* stands for *ACTIN2*. **C**, Comparison of the AZ enlargement and cell expansion in lines locally overexpressing WRKY57, IDA, and the previously published constitutive *p35S::IDA*. See also Supplemental Figure S9. Scale bars are 0.4 mm for the panels on the left and 20 µm in the confocal images. PI stands for propidium iodide, used to stain the cell walls (and some nuclei). GFP is shown in green and PI in grey. **D**, Example of cells that detach from the abscission zones of the *pHAE::WRKY57-GFP* lines while imaging. GFP fluorescence is shown in green, chlorophyll autofluorescence in magenta. Faint WRKY57-GFP signal can be observed in the cell periphery – most likely the cytoplasm – in samples that express the protein strongly. Scale bar represents 20 µm. **E**, Plants locally overexpressing *WRKYsrdx* in AZs strongly retain their floral organs. Scale bar represents 1 cm. **F**, Quantification of transgene expression levels in the transgenic lines shown in E. Same description as in B applies. **G**, Quantification of the floral organ abscission defect in the transgenic lines overexpressing *WRKY57srdx* in AZs. Markers (circle, square, triangle) represent the mean number of floral organs fully attached to the respective floral position. Whiskers are the SEM. White stars indicate statistically significant differences with wild-type Col-0 in one way ANOVA and post hoc Bonferroni tests at p < 0.05. Twelve plants per line were grown and one flower per plant and floral position was analyzed.

Conversely, we overexpressed in AZs a chimeric WRKY57 fused to the SRDX repressor domain. The addition of the SRDX repressor domain to the C-terminus of a TF drives transcriptional repression of its target genes (Hiratsu et al., 2003; Heyl et al., 2008; Matsui and Ohme-Takagi, 2010; Mahfouz et al., 2012; Cen et al., 2016). Multiple independent lines expressing *WRKY57srdx* under the *HAE* promoter showed mild to very strong retention of floral organs in developing siliques, showing a correlation between transgene expression level and phenotype severity (Figure 4E, F and G). Opposite phenotypic outcomes on abscission when locally overexpressing WRKY57 or WRKY57srdx indicate that this TF acts as a positive regulator at the developmental stage in which AZ cells are competent to abscise. We observed induction and repression of the *IDA* promoter activity in luciferase transient transactivation assays in Arabidopsis leaf mesophyll protoplasts (Supplemental Figure S10A). WRKY57 transactivation of *IDA* promoter was shown to be partially dependent on the presence of the five WRKY TFBSs disrupted in the *WRKY*(-) version of this promoter (Supplemental Figure S10B). These results suggest that WRKY57 positively regulates abscission by activating the *IDA* signaling pathway.

### Activation of abscission by WRKY57 requires IDA and redundant IDA-like peptides, as well as the receptors HAE and HSL2

Next, we explored the epistasis between the *pHAE::WRKY57-GFP* transgene, *IDA*, and *HAE/HSL2*. We crossed the *ida-2* and *hae hsl2* mutants to the single copy line *pHAE::WRKY57-GFP* (line 6), and obtained double homozygous *ida-2 pHAE::WRKY57-GFP* and triple homozygous *hae hsl2 pHAE::WRKY57-GFP* plants. Microscopic examination of the receptacle and AZ cells confirmed the effect of the local overexpression of *WRKY57* in AZs depends on *IDA* and the receptors *HAE* and *HSL2* (Figure 5A). Plants lacking functional receptors HAE/HSL2 completely suppressed the AZ enlargement and cell expansion phenotypes; however a weak but statistically significant enlargement was observed in the *ida-2 pHAE::WRKY57-GFP* double mutants (Figure 5B). We sought to independently confirm this result by transforming *ida-1* and *ida-2* alleles of *IDA* with the *pHAE::WRKY57-GFP* construct. Multiple independent lines displayed varying degrees of AZ enlargement and the enlargement correlated with a complementation of the abscission defect of *ida*, suggesting that induction of abscission by WRKY57 is only partially dependent on *IDA* while fully dependent on *HAE/HSL2* (Supplemental Figure S11A). We reasoned WRKY57 could also activate HAE/HSL2 via *IDL1*, *IDL2* or *IDL3* – close homologs of *IDA*. These three *IDL* genes contain several WRKY TFBSs in their promoter sequences, were reported to be expressed in floral receptacles, and expected to play a redundant role with *IDA* in abscission (Supplemental Figure S11B; (Vie et al., 2015)). Transcriptional upregulation of *IDA*, *IDL2* and *IDL3* was detected by RT-qPCR analysis from dissected floral receptacles in *pHAE::WRKY57-GFP* lines, and the capacity of WRKY57 to transactivate *IDL* genes was confirmed in transient luciferase assays in *Nicotiana benthamiana* leaves with the *IDL2* promoter (Figure 5C, Supplemental Figure S11C). These findings support a model where WRKY57 works as a positive regulator of abscission by orchestrating the coordinated expression of several redundant IDL peptides to activate the receptors HAE/HSL2.

**Figure 5.**
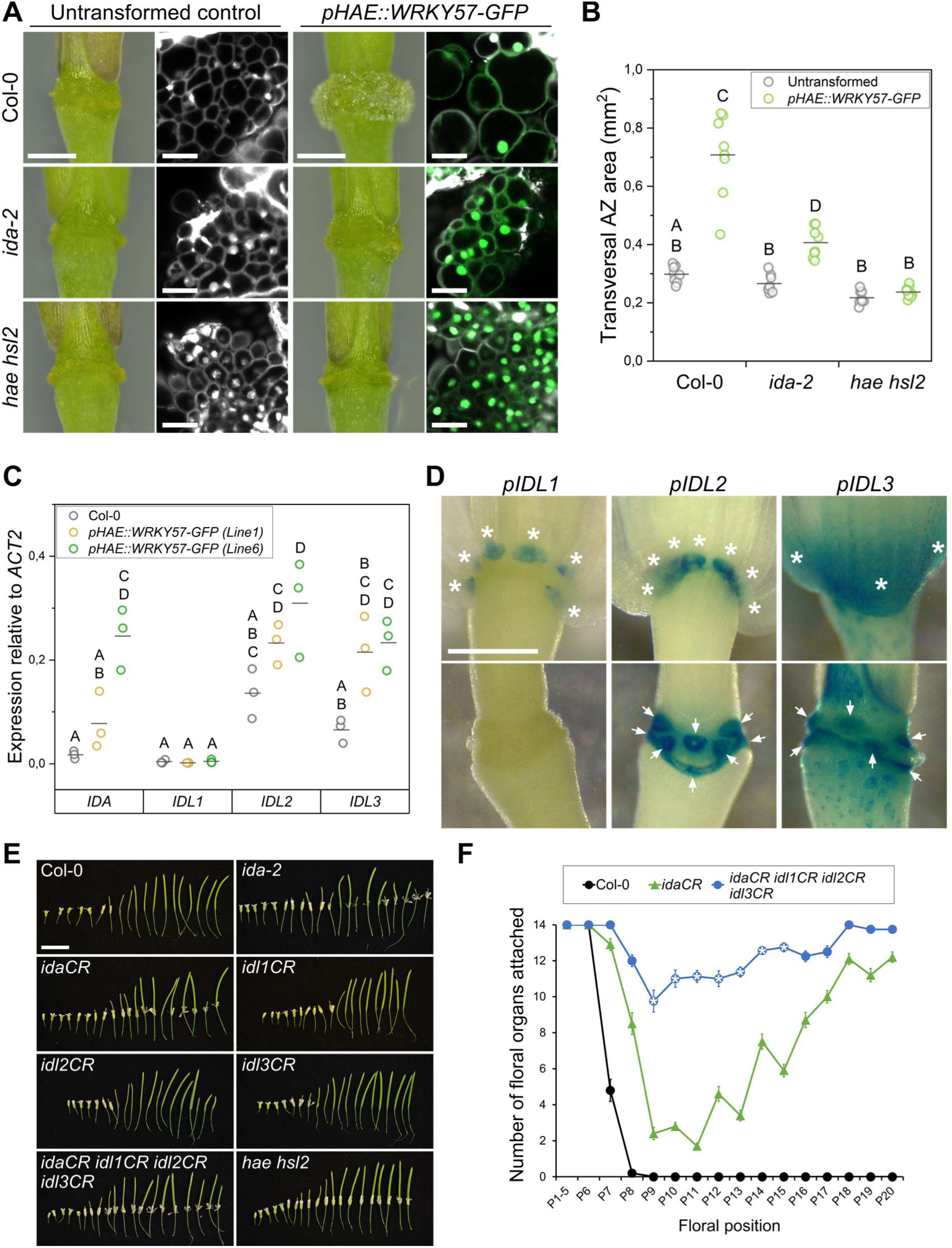
WRKY57 activates abscission via redundant *IDA* and *IDA*-*like* genes. **A**, Comparison of the AZ enlargement and cell expansion in lines locally overexpressing WRKY57-GFP in wild-type, *ida-2* and *hae hsl2* mutant backgrounds. See also Supplemental Figure S11. Scale bars are 0.4 mm for the panels on the left and 20 µm in the confocal images. GFP is shown in green and PI in grey. **B**, Quantification of the transversal AZ area in the genotypes displayed below the graph, either untransformed, or expressing the *pHAE::WRKY57-GFP* transgene. Data correspond to the measurement of the position 12 AZ area in the main inflorescence stem for each plant. Eight plants per genotype were analyzed. Letters show statistically significant differences between groups in one-way ANOVA and Bonferroni post hoc analyses (p < 0.05). **C**, Estimation of gene expression of *IDA*, *IDL1*, *IDL2* and *IDL3* in the genotypes listed in the graph legend. Data correspond to the average of two technical replicates per biological replica. Three biological replicas were assayed per genotype. Letters show statistically significant differences between groups in two-way ANOVA and Bonferroni post hoc analyses (p < 0.05). **D**, Histochemical detection of GUS activity in plants expressing GFP-GUS under the control of the promoters listed over the panels. White stars highlight the floral organs still attached in the upper panels. White arrows point at the residuum cells with clear GUS activity in the receptacle. Scale bar represents 0.4 mm. **E**, Floral organ abscission in the genotypes listed in the panels. Scale bar represents 1 cm. **F**, Quantification of retained floral organs per floral position along the main inflorescence stem for the genotypes listed in the graph’s legend. Markers (circles and triangles) represent the mean; whiskers are the SEM. White stars denote the floral positions of the quadruple *idaCR idl1CR idl2CR idl3CR* that were significantly different from *idaCR* in one way ANOVA and Bonferroni post hoc tests (p < 0.05). See Supplemental Figure S12 for the expanded quantification of all genotypes included in panel E.

The abscission inducing activity of *IDL1*, *IDL2* and *IDL3* was demonstrated in constitutive overexpression lines (Stenvik et al., 2008). IDL2 and IDL3 were shown to bind with high affinity the complex formed by HAE and SERK extracellular domains, as well as the extracellular domain of HSL1 – closest homolog to HAE/HSL2 in Arabidopsis (Roman et al., 2022). Genetic evidence involving *IDL1*, *IDL2* and *IDL3* in floral organ abscission has nevertheless remained elusive. Promoter reporter lines for *IDL1*, *IDL2* and *IDL3* expressing GUS-GFP were generated and their previously reported expression patterns during abscission confirmed (Stenvik et al., 2008; Vie et al., 2015). *IDL1* promoter activity was detected in the secession cells only, while in the case of *IDL2* and *IDL3*, promoter activities were detected in both residuum and secession cell layers (Figure 5D). Gene-edited knock-out lines for *IDL2* and *IDL3* were generated in the Col-0 background with the CRISPR-Cas9 technology, and higher order mutants were obtained by genetic crosses with previously published *idaCR* and *idl1CR* (Shi et al., 2018; Alling and Galindo-Trigo, 2023). The *idaCR* line was used despite the availability of the *ida-2* allele, also in Col-0, due to T-DNA-induced genomic rearrangements in the *ida-2* line that impede obtaining double mutants between *ida* and *idl1* alleles (Alling and Galindo-Trigo, 2023). Developing siliques in the quadruple mutant *idaCR idl1CR idl2CR idl3CR* retained floral organs more strongly than the single *ida* mutant, confirming the genetic redundancy between these IDL peptides as positive regulators of abscission (Figure 5E, F and Supplemental Figure S12). Based on the cumulative evidence provided here, we propose a model in which WRKY57 modulates the transcription of *IDA* and *IDL* genes to fine-tune the timing of floral organ abscission.

## Discussion

The multiplicity of cues affecting abscission and the expansion of TF families in flowering plants make characterizing transcriptional effectors in this process a demanding task. In this study, we show that the transcriptional regulation of *IDA*, one of the main triggers of floral organ abscission, is subject to the control of TFs from the ERF and WRKY families (Figure 1). Using molecular genetics and physiological assays, we showed that the IDA signaling pathway can be activated upon exposure to a bacterial immune elicitor in AZs, and that this activation depends on WRKY TFBSs in the *IDA* promoter (Figure 2). Screening a large collection of TFs in a heterologous system, we identified a list of potential candidates to directly regulate *IDA* expression, and genetically characterized WRKY57 as such (Figure 3, 4 and 5A and B). Finally, by locally overexpressing WRKY57 in AZs we found evidence of coordinated transcriptional regulation of *IDA*, *IDL2* and *IDL3* (Figure 5C). The generation of higher order *ida/idl* mutants confirmed the long-standing hypothesis of functional redundancy among these genes (Figure 5E and F).

ERFs are known to relay the transcriptional signaling induced by ethylene downstream of the master transcriptional regulator ETHYLENE INSENSITIVE3 (EIN3; (Chao et al., 1997; Chang et al., 2013)). Indeed, the result of disrupting one ERF TFBSs in the *IDA* promoter yielded the same expression pattern observed for the wild-type *IDA* promoter in an ethylene insensitive background (Butenko et al., 2006). It is therefore plausible that ethylene has a lead role in activating *IDA* transcription during the developmentally initiated abscission pathway. Especially so when considering the lack of genetic complementation of the abscission phenotype of *ida* mutant plants when *IDA* expression is driven by a promoter that lacks ERF TFBSs (Figure 1C). Interestingly, the master regulator of ethylene EIN3 was identified in the Y1H screening as a TF that binds the *IDA* promoter (Supplemental Table S3). EIN3 upregulates the transcription of its direct targets, providing further support to the model in which *IDA* is activated by ethylene signaling during floral organ abscission (Chang et al., 2013; Meir et al., 2019). Although *IDA* was not detected in the genome-wide screen for EIN3-mediated, ethylene-induced transcriptionally regulated genes in Arabidopsis, developmental or tissue-specific effects could explain its absence, as these experiments were conducted in 3-day old etiolated seedlings (Chang et al., 2013). Dissecting the importance and the mechanism behind the putative two-tiered regulation of *IDA* by EIN3 and ERFs will be the topic of future investigations.

We have shown that residuum cells of recently abscised floral receptacles (positions six or seven) respond to flg22 treatments by upregulating *IDA* (Figure 2A and B). While the fluorescent reporter data suggests residuum cells are particularly responsive to flg22, this responsiveness pattern may not reflect responsivity *per se*, but rather their capacity to uptake the treatment. Residuum cells synthesize a cuticle after abscission takes place, although in the floral positions analyzed here this process may not have concluded, presenting a less hydrophobic exterior and increased permeability. Regardless of the spatial specificity of the response, the responsivity of the *IDA* promoter in residuum cells to biotic stress after organs abscise has interesting implications. Firstly, it reinforces the notion that IDA signaling is activated in response to flg22 in cells destined for separation. Secondly, reactivation of IDA signaling in the cells that control cell separation will likely incur in the activation of a similar set of responses triggered during abscission, as the expression of receptors HAE/HSL2 and other downstream regulators remains active in residuum cells after separation (Cho et al., 2008). Flg22 elicitation of *IDA* expression could induce a protective shedding mechanism that is useful for the plant when still-attached floral organs are infected by bacterial pathogens, similar to pathogen-induced cauline leaf shedding (Patharkar et al., 2017). Alternatively, activating IDA signaling could help residuum cells protect themselves against pathogens by reinforcing the flg22-mediated signaling. When seedlings are exposed to exogenous IDA peptide, defense response marker genes that are typically induced by flg22 like *FLG22-INDUCED RECEPTOR-LIKE KINASE1* (*FRK1;* (Asai et al., 2002)) are transcriptionally upregulated. When seedlings are exposed to both IDA and flg22, the expression of these marker genes is upregulated even further (Lalun et al., 2023). This suggests a role for IDA in enhancing the flg22-mediated transcriptional response, something potentially beneficial for residuum cells to protect them as cell separation occurs. There is however conflicting evidence that indicates a negative role during leaf bacterial colonization for the IDA receptors HAE/HSL2 and the related peptide IDL6 (Wang et al., 2017). Understanding the physiological consequence of the molecular signatures triggered by IDA treatments that overlap with immune responses awaits further analyses. AZ-centric physiological studies will provide exciting new insights into the relationship between abscission and immunity.

Out of the twelve WRKYs found in the Y1H screen on the *IDA* promoter, there are several good candidates to regulate *IDA* during biotic stress. In ChIP-seq and mass spectrometry analyses of nuclear proteins, Birkenbihl et al. reported the *IDA* promoter-binding TFs WRKY8 and WRKY11 to be both transcriptionally upregulated and to increase in protein abundance upon flg22 treatments (Birkenbihl et al., 2018). WRKY8 was found to execute most of the transcriptional responses in chitin-induced, PBS1-LIKE 19 (PBL19)-mediated immunity. Upon chitin treatments – also known to induce *IDA* expression –, PBL19 accumulates in the nucleus, interacts and phosphorylates WRKY8 which in turn transcriptionally upregulates genes. Transgenic plants expressing a version of PBL19 that constitutively accumulates in the nucleus show a severe autoimmunity phenotype, which is dependent on WRKY8. An RNAseq analysis of the autoimmune PBL19 expressing plants revealed that *HAE*, *IDL6* and *IDL7* were upregulated along with immunity marker genes like *FRK1* – data for the *IDA* gene was unfortunately not available in the dataset (Li et al., 2022). This suggests that upon chitin perception, WRKY8 induces expression of components of the IDA signaling pathway, and quite possibly also *IDA*. Since WRKY8 increases in abundance upon flg22 detection, it is likely that a similar set of genes are upregulated by WRKY8 in the context of flg22 perception; making WRKY8 a prime candidate to mediate the flg22-induced IDA expression. WRKY11, on the other hand, acts as a negative regulator of gene expression and inhibits transcription of genes typically induced in the context of immunity redundantly with WRKY7, WRKY15, WRKY17, WRKY21 and WRKY39. Quintuple mutants *wrky7-11-17-21-39* show severe developmental defects and display constitutive upregulation of defense-related genes. An exploration of the RNAseq dataset comparing *wrky7-11-17-21-39* to wild-type confirmed that the IDA signaling pathway is also differentially expressed, with *IDA*, *HAE*, *IDL6* and *IDL7* being upregulated in the quintuple *wrky* mutant (Du et al., 2023). This set of WRKYs likely constitute a negative feedback loop to the defense-induced transcriptional responses in the plant, including components of the IDA signaling pathway.

We have genetically characterized WRKY57 as a potential positive regulator of abscission, exerting its transcriptional control on *IDA*, *IDL2* and *IDL3* (Figure 4 and 5). In rosette leaves, WRKY57 can be co-opted by repressors from the jasmonic acid and auxin signaling pathways to respectively induce or repress jasmonic acid-induced senescence (Jiang et al., 2014). Both hormones have important roles in the abscission and senescence of floral organs. While auxin inhibits abscission (Basu et al., 2013), jasmonic acid signaling regulates floral organ senescence and the timing of abscission (Kubigsteltig et al., 1999; Serrano-Bueno et al., 2022). Thus, we speculate that WRKY57 could help orchestrate the coordinated transcriptional regulation of the *IDA* gene from auxin and jasmonic acid signaling pathways to fine tune abscission timing.

The regulatory axis described in this study comprising WRKY-IDA signaling-abscission is not exclusive to floral organ abscission in Arabidopsis. Tomato pedicels have been recently shown to utilize a WRKY TF to mediate low light-induced pedicel abscission by regulating tomato *IDL6* gene expression (Li et al., 2021). This demonstrates the tight relationship between transcriptional regulators of the WRKY family and abscission is conserved across plant species and in diverse plant organs.

Introducing beneficial traits into crops by targeted genome editing is easier than ever; however, gene functions are normally multifaceted and involved in multiple processes. Our study exemplifies how promoter element modification can be exploited to allow the developmentally encoded expression profile of a gene to proceed, while impairing its responsiveness to stress (Figure 1 and 2). Promoter targeted genome editing approaches therefore show promise in allowing the introduction of beneficial traits like controlled abscission into crops, while minimizing developmental penalties to yield (Liu et al., 2021; Tang and Zhang, 2023; Zhou et al., 2023).

## Abbreviations

AZ: Abscission zone
ERF: Ethylene response factor
HAE: HAESA
IDA: INFLORESCENCE DEFICIENT IN ABSCISSION
IDL: IDA-like
MAMP: Microbe-Associated Molecular Pattern
TF: Transcription factor
TFBS: TF binding sites
Y1H: Yeast one-hybrid

## Supplementary data

**Supplemental Table S1.** Primers used in this study.

**Supplemental Table S2.** Transcription factor binding site prediction for the *IDA* promoter.

**Supplemental Table S3.** Transcription factors identified to bind the *IDA* promoter in the yeast one-hybrid screening.

**Supplemental Figure S1.** Transcription factor binding site analysis of the *IDA* promoter sequence in Arabidopsis.

**Supplemental Figure S2.** Additional information to main Figure 1B.

**Supplemental Figure S3.** Histochemical detection of GUS activity during lateral root emergence.

**Supplemental Figure S4.** The *ERF(-)* version of the *IDA* promoter cannot rescue the abscission defect of the *ida-1* mutant.

**Supplemental Figure S5.** Deletion of individual WRKY binding sites in the *IDA* promoter does not impair its activity during abscission.

**Supplemental Figure S6.** Elicitation of immune responses in cauline leaves with flg22 does not induce IDA in the cauline AZ.

**Supplemental Figure S7.** Additional information to main Figure 3.

**Supplemental Figure S8.** Nuclear localization of the fusion proteins WRKY60-GFP and WRKY57-GFP.

**Supplemental Figure S9.** Quantification of the receptacle enlargement phenotype shown in the main Figure 4.

**Supplemental Figure S10.** Transient transactivation of the *IDA* promoter by WRKY57 in Arabidopsis mesophyll protoplasts.

**Supplemental Figure S11.** WRKY57 activates the abscission pathway in *ida* mutants via *IDL* peptides.

**Supplemental Figure S12.** Quantification of floral organ abscission at positions 8, 10, 12, 16 and 20.

## Acknowledgements

The authors thank Prof. Dr. Reidunn Aallen for sharing the *idl1CR*, *idl2CR* and *idl3CR* lines; Prof. Dr. Rüdiger Simon for sharing the H2B-TdTomato plasmid; and Verena Mertes for her initial contributions to the Y1H screening. We thank the NorMic microscopy platform for their help and support. We thank the technician team at EVOGENE for their technical support.

## Author contributions

S.G-T. conceived and developed the research hypotheses, produced the genetic materials, carried out the experimentation, data analyses and interpretation, wrote the manuscript and prepared all figures and illustrations. T.I. and S.S. provided the *idl2CR* and *idl3CR* lines. A-M.B. cloned the Y1H constructs and carried out the Y1H screen together with S.B. M.A.B. designed the Y1H screen of the *IDA* promoter with S.B., acquired funding, administered resources, and provided initial guidance interpreting the abscission phenotypes. All authors read the manuscript and provided feedback.

## Data availability

All data presented can be found in the manuscript and supplementary data. Requests for further details can be directed to the corresponding authors.

