## Supplemental_Figures for "ERF and WRKY transcription factors regulate *IDA* and abscission timing in Arabidopsis"

**Supplemental Figures (Galindo-Trigo et al. 2023)**

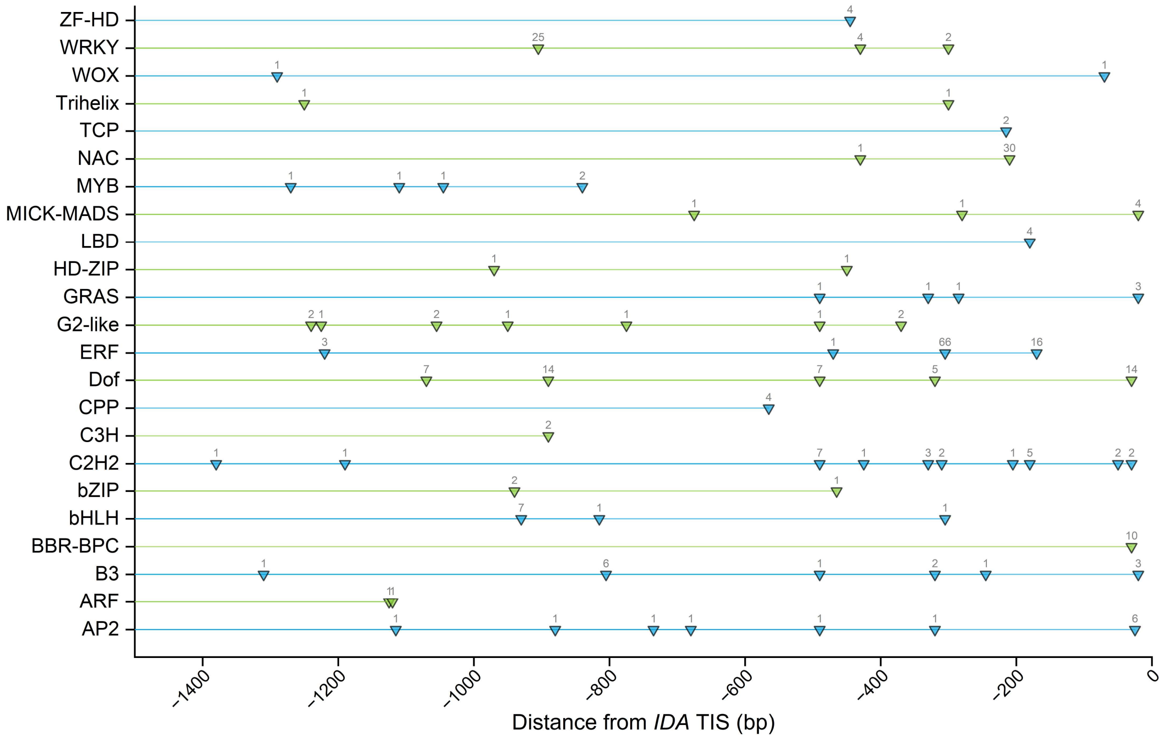


**Supplemental Figure S1.** Transcription factor binding site analysis of the *IDA* promoter sequence in Arabidopsis. Representation of the presence and location of each of the TFBSs predicted for each of the main TF families in Arabidopsis using the Binding Site Prediction tool from PlantRegMap. Inverted triangles along each line represent the predicted TFBSs for its respective TF type – listed on the y axis. The number over each inverted triangle indicates the number of individual TFs that were predicted to bind at that specific binding site. TIS, translational initiation site.

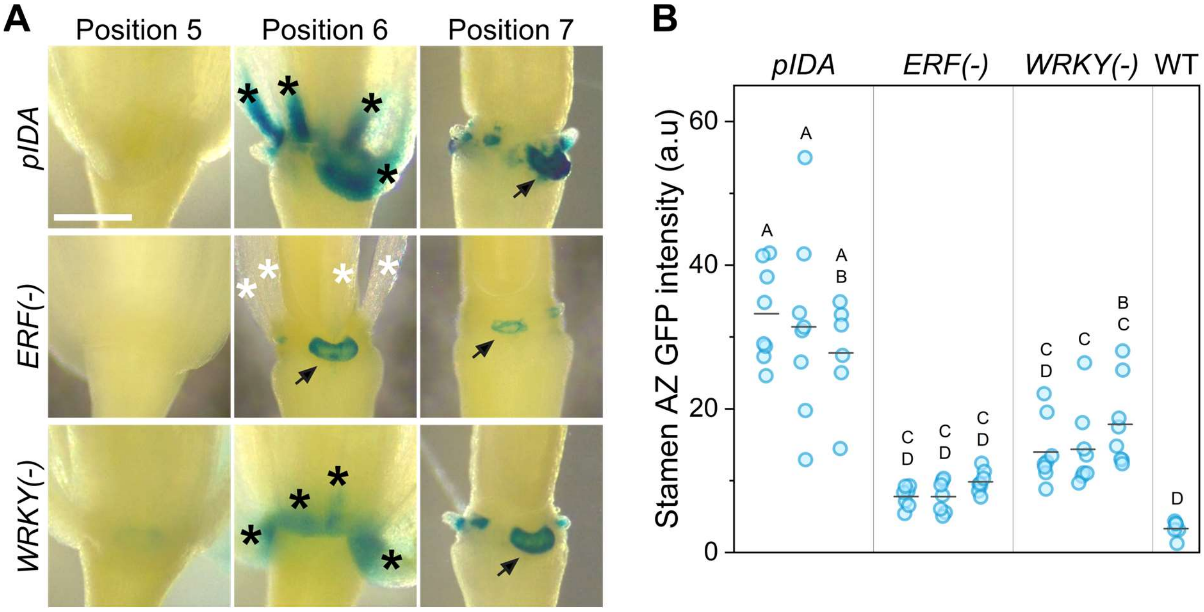


**Supplemental Figure S2.** Additional information to main Figure 1B. **A,** Histochemical detection of beta-glucuronidase (GUS) activity in flowers at the indicated floral positions. The GFP-GUS gene was driven by the promoters indicated on the left. Black stars indicate floral organs with intense GUS activity. White stars denote floral organs devoid of GUS activity. Arrows point at the nectaries. Scale bar is 0.4 mm. **B,** Quantification of the GFP fluorescence intensity in the abscission zone cells of stamens from position 6 flowers in transgenic plants where GFP-GUS expression was driven by the promoters indicated on top of the graph. One flower per plant and eight plants were assayed per transgenic line. Each section of the graph contains the data from three independent transgenic lines per construct. Different letters denote statistically significant differences in two-way ANOVA analyses and post hoc pairwise Bonferroni tests ($p < 0.05$). A.U. stands for arbitrary units.

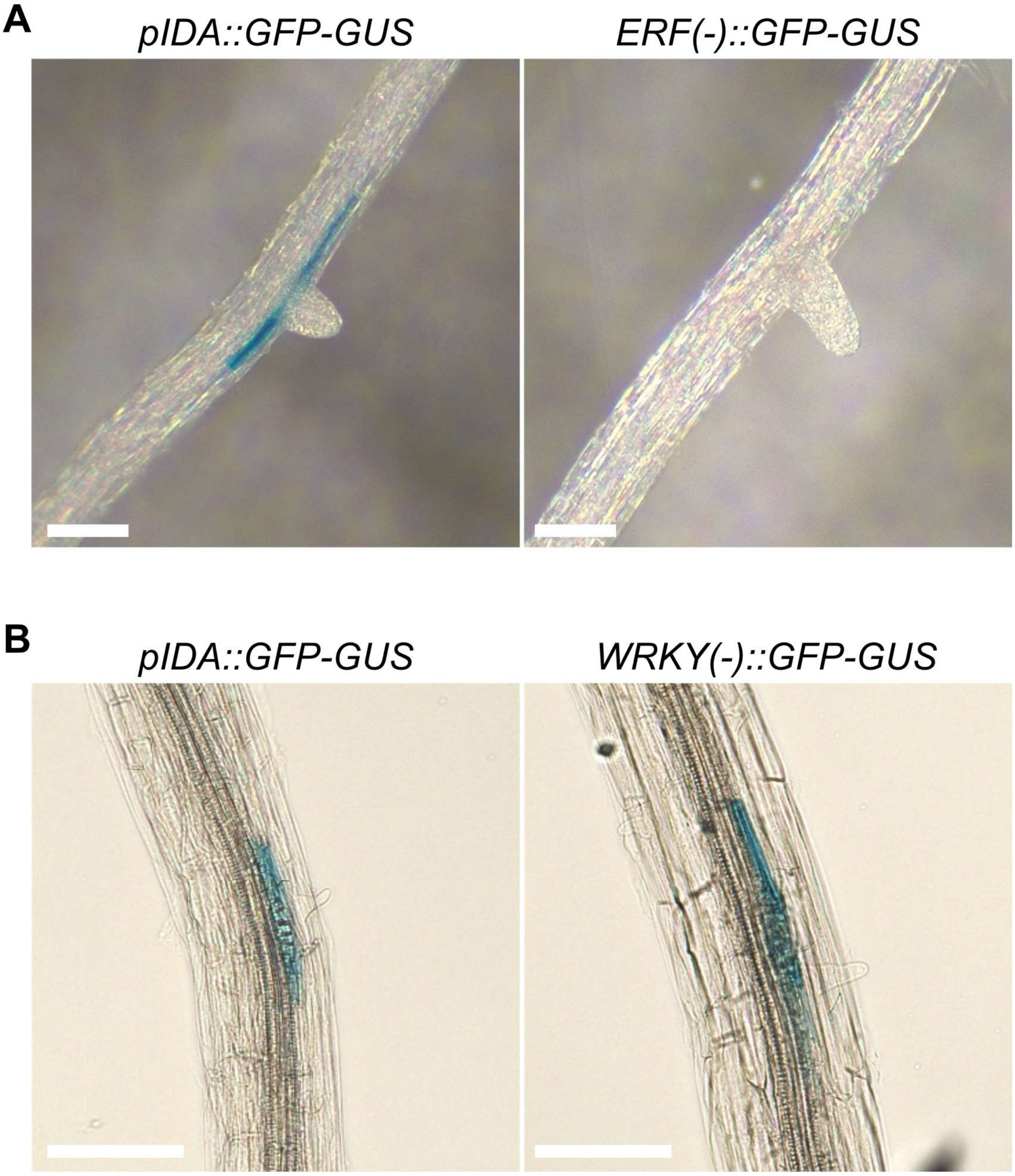


**Supplemental Figure S3.** Histochemical detection of GUS activity during lateral root emergence. **A,** The *ERF*(-) version of the *IDA* promoter is not active during lateral root emergence in ten-day-old seedlings. **B,** The *WRKY*(-) version of the *IDA* promoter is still active during lateral root emergence. Pictures are representative of three or more independent transgenic lines for each of the constructs. All constructs were transformed into Col-0 plants. Scale bars are 50 µm.

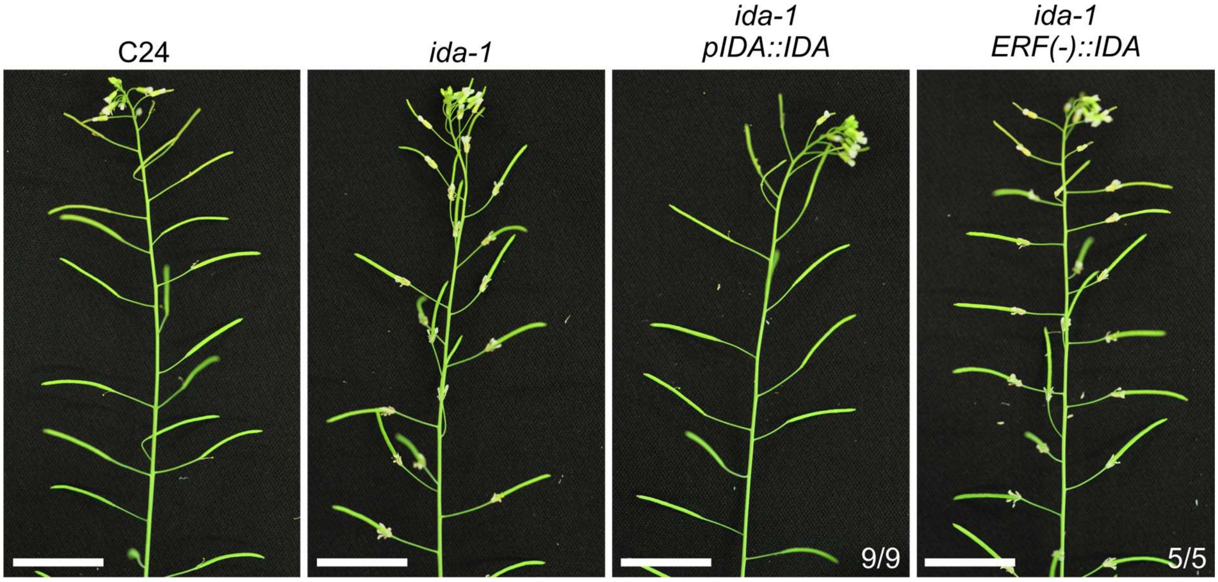


**Supplemental Figure S4.** The *ERF(-)* version of the *IDA* promoter cannot rescue the abscission defect of the *ida-1* mutant. Representative pictures of a transgenic line per construct are shown. Numbers indicate the number of independent transgenic lines obtained displaying the phenotype shown in the picture. Scale bars are 2 cm.

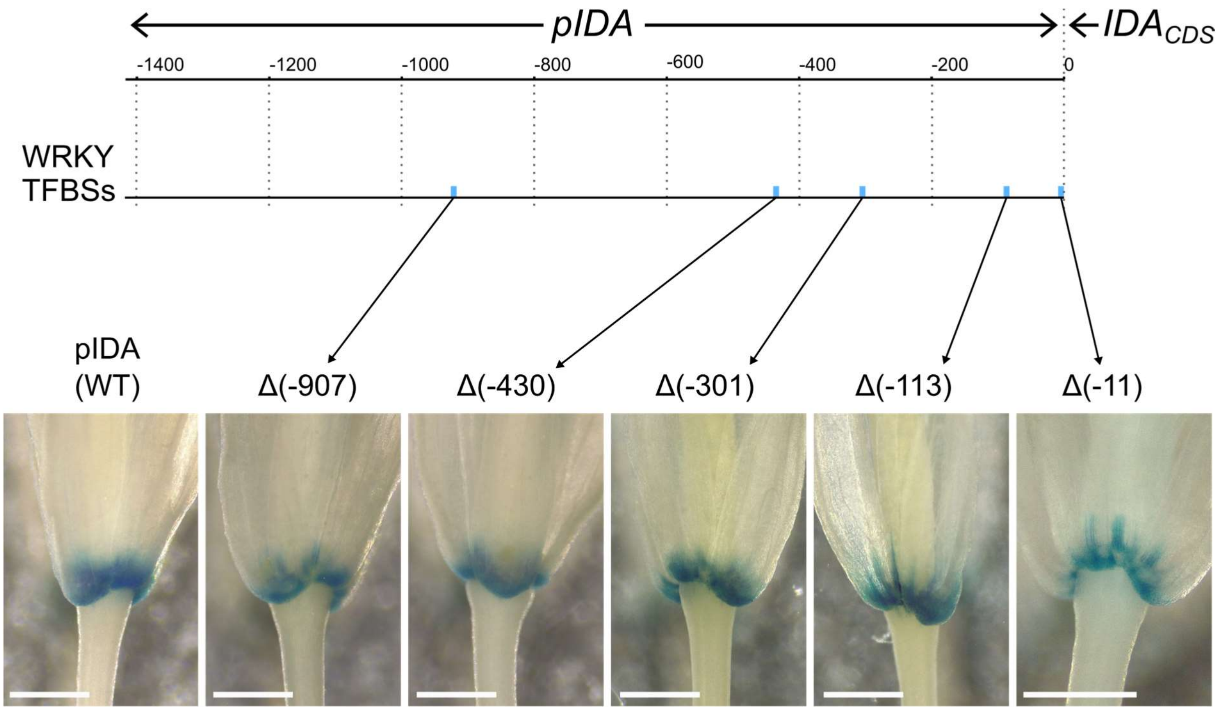


**Supplemental Figure S5.** Deletion of individual WRKY binding sites in the *IDA* promoter does not impair its activity during abscission. The upper diagram represents the *IDA* promoter sequence and the position of the predicted WRKY transcription factor binding sites (TFBSs; blue rectangles). Below are representative images of histochemical detection of GUS activity in reporter lines expressing GFP-GUS under the control of the wild-type *IDA* promoter, or versions of the *IDA* promoter in which each of the five WRKY TFBSs were disrupted by site-directed mutagenesis. In brackets are the distance from the translational initiation site of *IDA*. The images shown here are representative of three or more independent transgenic lines obtained for each construct. GUS activity detection assays were not conducted in parallel and therefore small differences in the relative intensity of the blue depositions should not be overinterpreted. Scale bars are 500 µm.

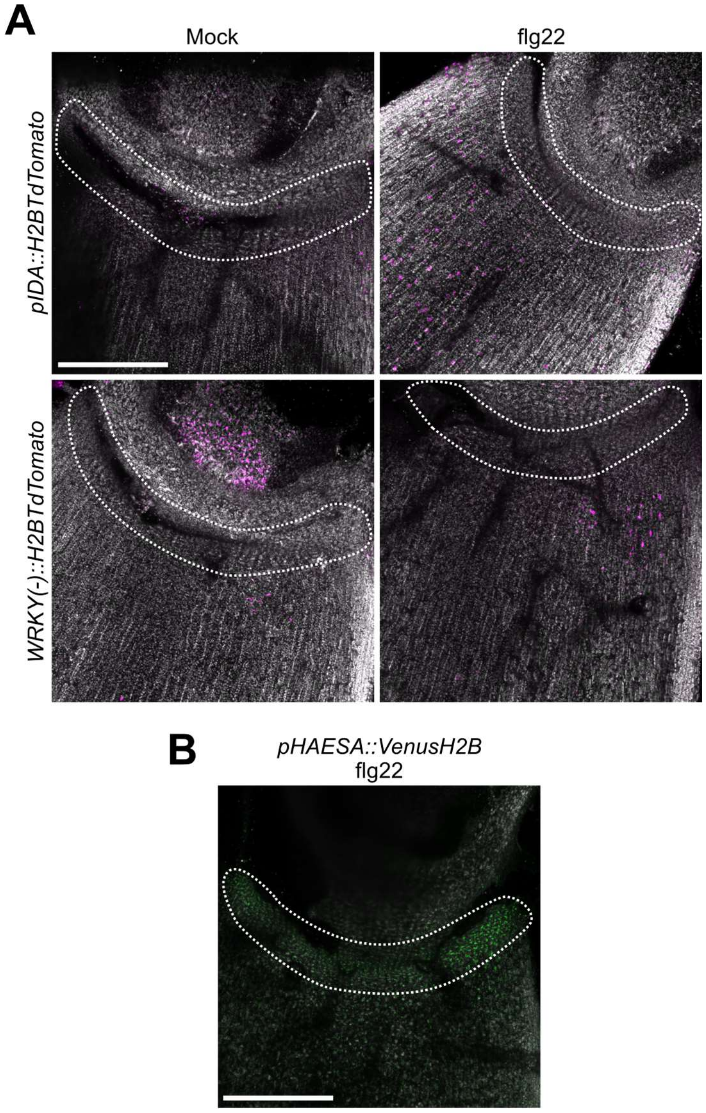


**Supplemental Figure S6.** Elicitation of immune responses in cauline leaves with flg22 does not induce *IDA* in the cauline AZ. **A,** Representative confocal micrographs of the experiment comparing *IDA* and *WRKY*(-) promoters 20h after treatment. Promoter activity is reflected by the presence of H2B-TdTomato tagged nuclei in the images. H2B-TdTomato is shown in magenta; Chlorophyll autofluorescence is shown in grey. Dotted lines surround the cauline leaf-inflorescence stem junction. The same results were seen in three independent lines per construct, assaying one cauline leaf per plant and five plants per construct and treatment combination. **B,** The *HAESA* promoter is active in the cauline leaf abscission zones after flg22 infiltration of the cauline leaves, as reported by Patharkar et al. (Patharkar et al., 2017).

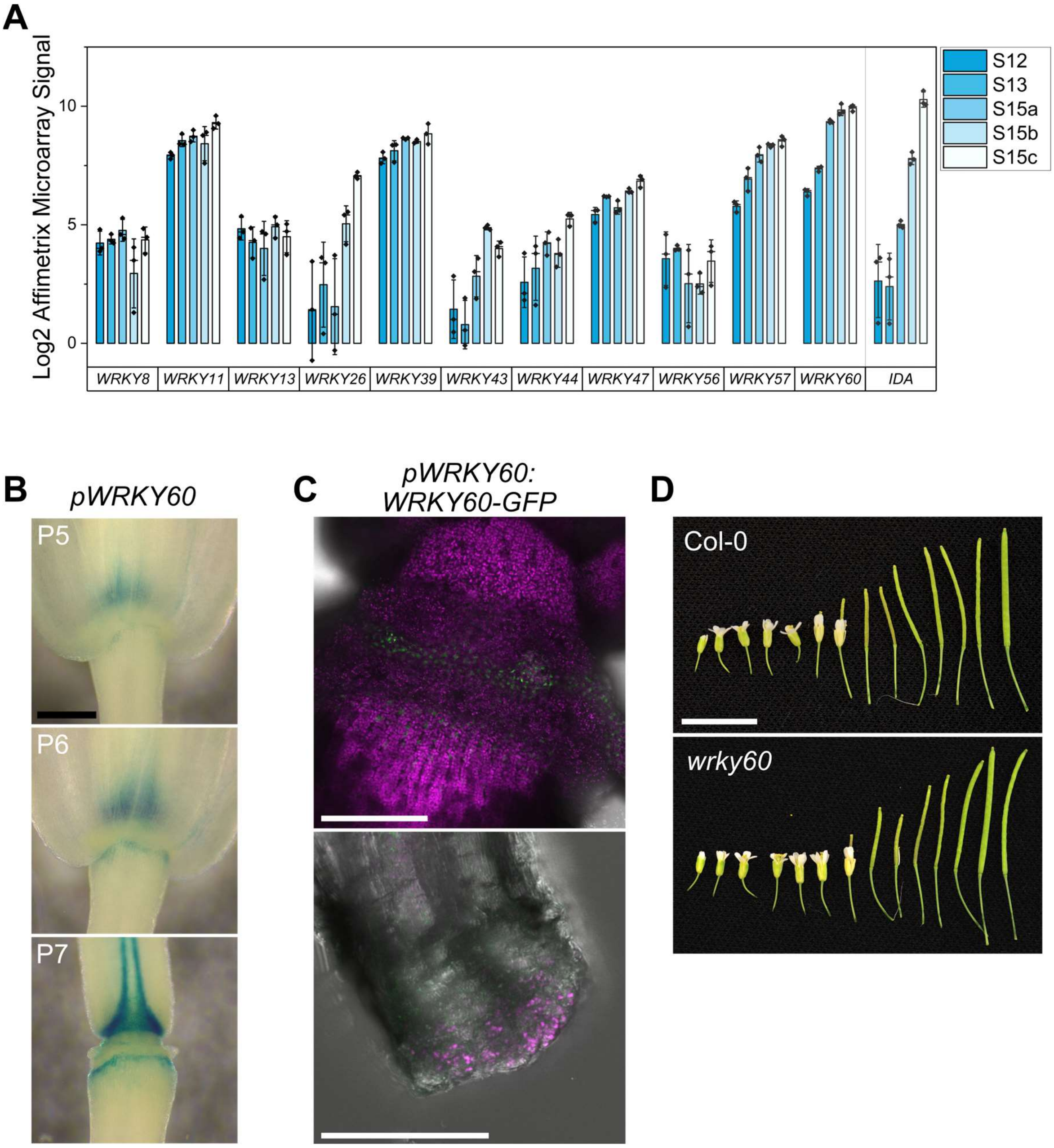


**Supplemental Figure S7.** Additional information to main Figure 3. **A,** Expression in stamen AZ cells in the floral stages S12 to S15c, which typically coincide with floral positions four to six in which abscission is initiated and takes place. *IDA* is also shown on the right. **B,** Histochemical staining of GUS activity in plants expressing GFP-GUS under the *WRKY60* promoter. Expression appears to preferentially take place outside of AZs. Scale bar is 0.4 mm. P5, P6, P7 are floral positions five, six and seven along the main inflorescence. **C,** The WRKY60-GFP protein is expressed and accumulates outside of abscission zones in the neighboring cells. Confocal micrographs showing GFP fluorescence in green and chlorophyll autofluorescence in magenta. Top picture shows the floral receptacle with the residuum abscission zone cells. The secession cells of a stamen are shown in the lower panel. Scale bars are 150 µm in the top image and 50 µm in the bottom picture. **D,** The single T-DNA insertional mutant *wrky60* does not present floral organ abscission defects. Scale bar is 1 cm.

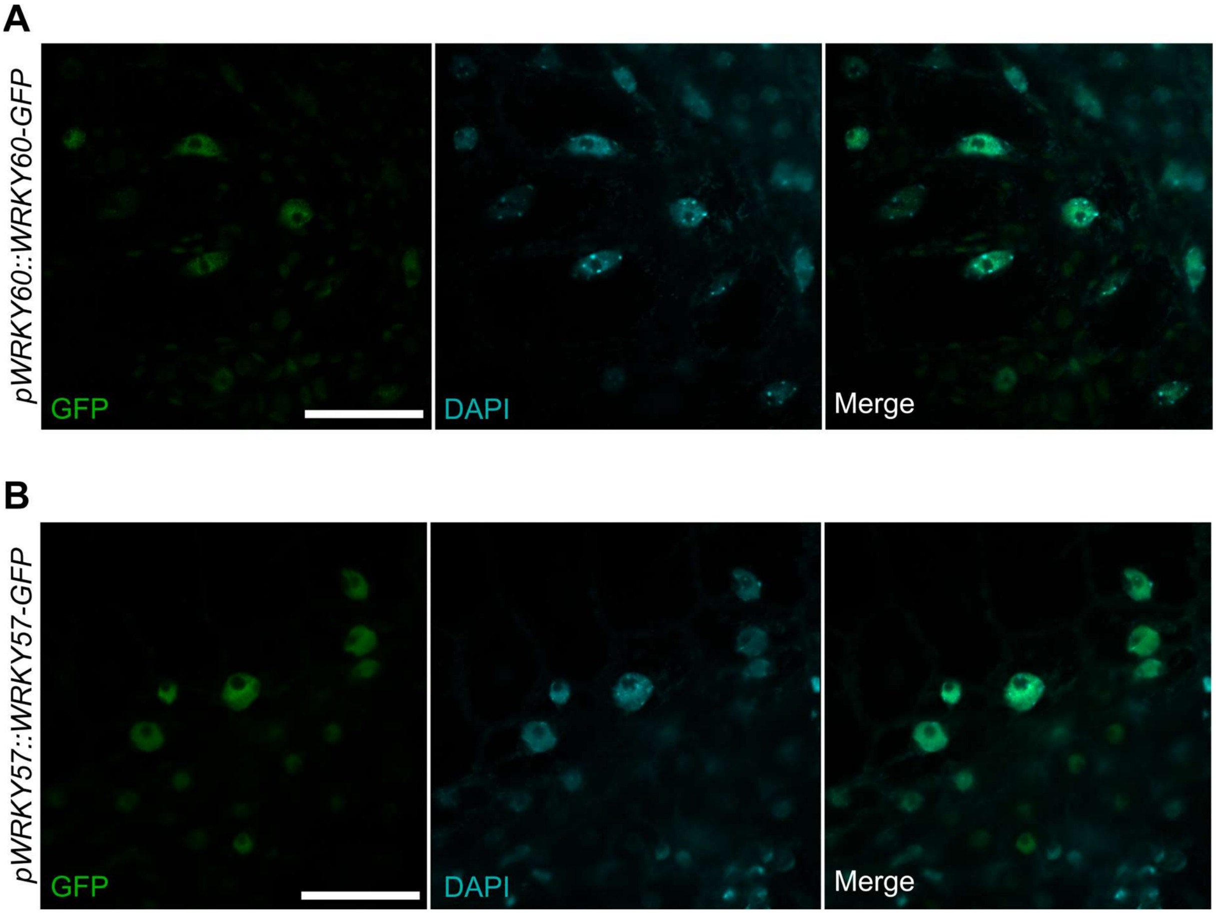


**Supplemental Figure S8.** Nuclear localization of the fusion proteins WRKY60-GFP and WRKY57-GFP. **A,** Nuclei from receptacle cells adjacent to AZs showing WRKY60-GFP accumulation in the nucleus (in green) and DNA (in blue, DAPI stained). **B,** Residuum cells from the receptacle AZ showing nuclear accumulation of WRKY57-GFP (in green), and DNA (in blue, DAPI stained). Scale bars are 25 µm in A and B.

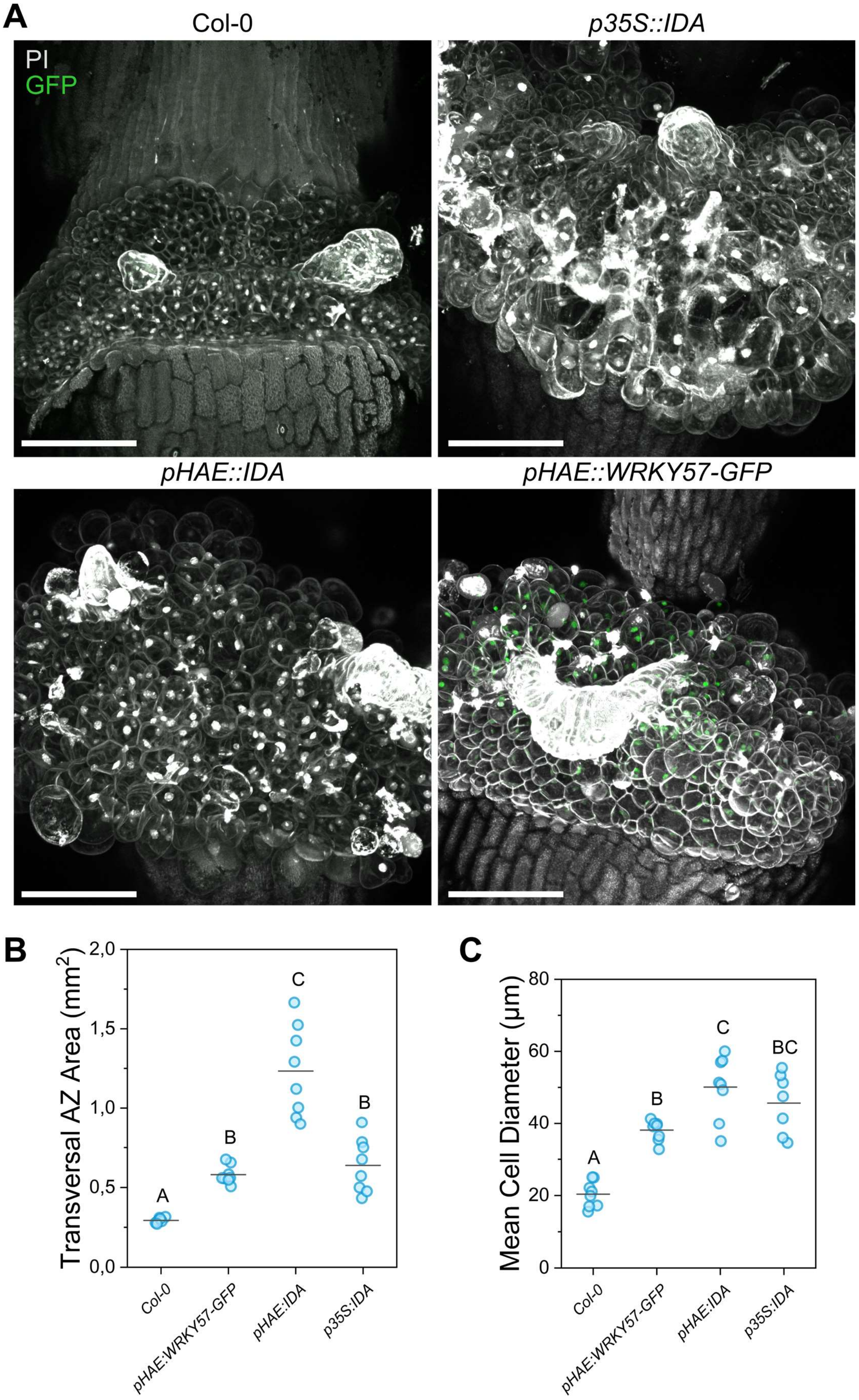


**Supplemental Figure S9.** Quantification of the receptacle enlargement phenotype shown in the main Figure 4. **A,** Maximum intensity projections of the floral receptacle of the genotypes indicated above the panels. Propidium iodide staining of cells walls (and some nuclei) is shown in grey. GFP fluorescence is shown in green. Scale bars are 100 µm. **B,** Quantification of the AZ enlargement phenotype shown in main Figure 4. Each datapoint corresponds to the transversal AZ area of the floral position 12 in the main inflorescence of a plant. Eight plants per genotype were analyzed. **C,** Quantification of the mean cell diameter of the genotypes listed below the graph. mean cell diameter was calculated for one flower per plant and eight plants per genotype. Letters in panels B and C represent statistically significant differences between groups in one way ANOVA and Bonferroni post hoc tests ($p < 0.05$).

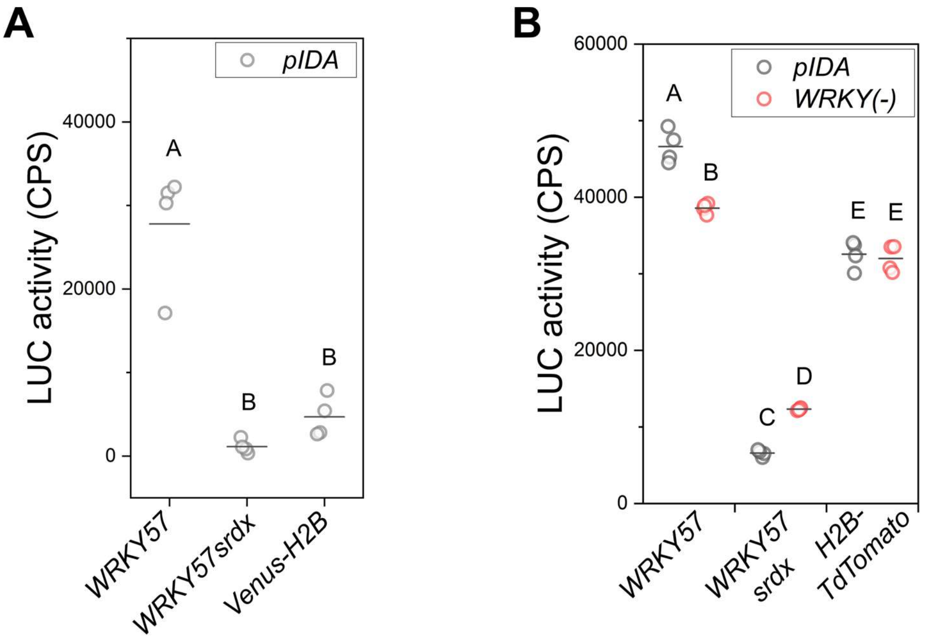


**Supplemental Figure S10.** Transient transactivation of the *IDA* promoter by WRKY57 in Arabidopsis mesophyll protoplasts. **A,** Luciferase activity in protoplasts co-transfected with *pIDA::LUC* and the effectors listed below the graph. *WRKY57*, *WRKY57srdx* and *Venus-H2B* were expressed under the constitutive promoter *p35S*. **B,** Comparison between the capacity of WRKY57 to transactivate the *IDA* promoter with and without the five WRKY TFBSs disrupted in *WRKY*(-). Data in both panels represent the light emission in counts per second (CPS) in four independent protoplast co-transfections. The experiments were repeated three times and similar trends were observed. Letters represent statistically significant differences between groups in one way ANOVA and post hoc Bonferroni pairwise comparisons ($p < 0.05$). Venus-H2B and H2B-TdTomato were used as negative controls to compare the effect of the overexpression of an unrelated nuclear protein in *IDA* promoter activity.

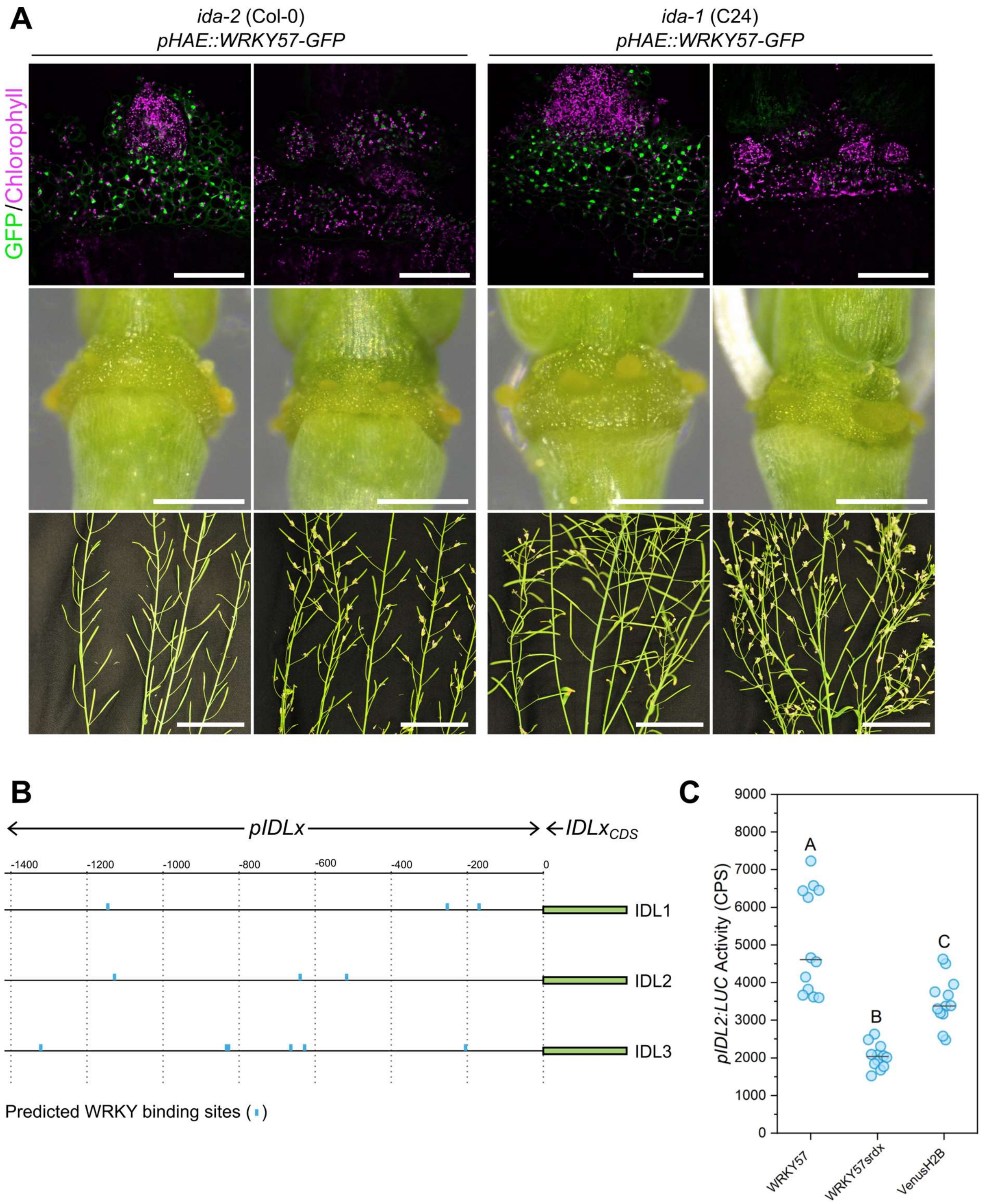


**Supplemental Figure S11**. WRKY57 activates the abscission pathway in *ida* mutants via *IDL* peptides. **A,** Representative examples of independent transgenic lines in which *pHAE::WRKY57-GFP* was transformed into *ida-2* or *ida-1*. For each genotype, two independent T1 plants with high and very low transgene expression levels are shown in different columns. For each line and from top to bottom, panels represent a confocal micrograph showing GFP fluorescence (green) and chlorophyll autofluorescence (magenta) in the receptacle AZs, a closeup of the AZ imaged in a stereomicroscope, and a picture of their floral organ abscission defects in mature plants. Scale bars are 100 µm, 300 µm, and 4 cm, respectively. The protein expression level, AZ size and complementation of the *ida* abscission phenotypes correlate. **B,** diagram of the presence of WRKY TFBSs in the promoter sequences of *IDL1*, *IDL2* and *IDL3*. **C,** Luciferase activity in transiently transformed *N. benthamiana* leaves with *pIDA::LUC* and the effectors listed below the graph. *WRKY57*, *WRKY57srdx* and *Venus-H2B* were expressed under the constitutive promoter *p35S*. Datapoints are the light emission recorded in counts per second (CPS) for 12 individual leaf discs. Letters show statistically significant differences between groups in a one way ANOVA and Bonferroni post hoc pairwise comparisons ($p < 0.05$). The luciferase assay was repeated twice with the same result.

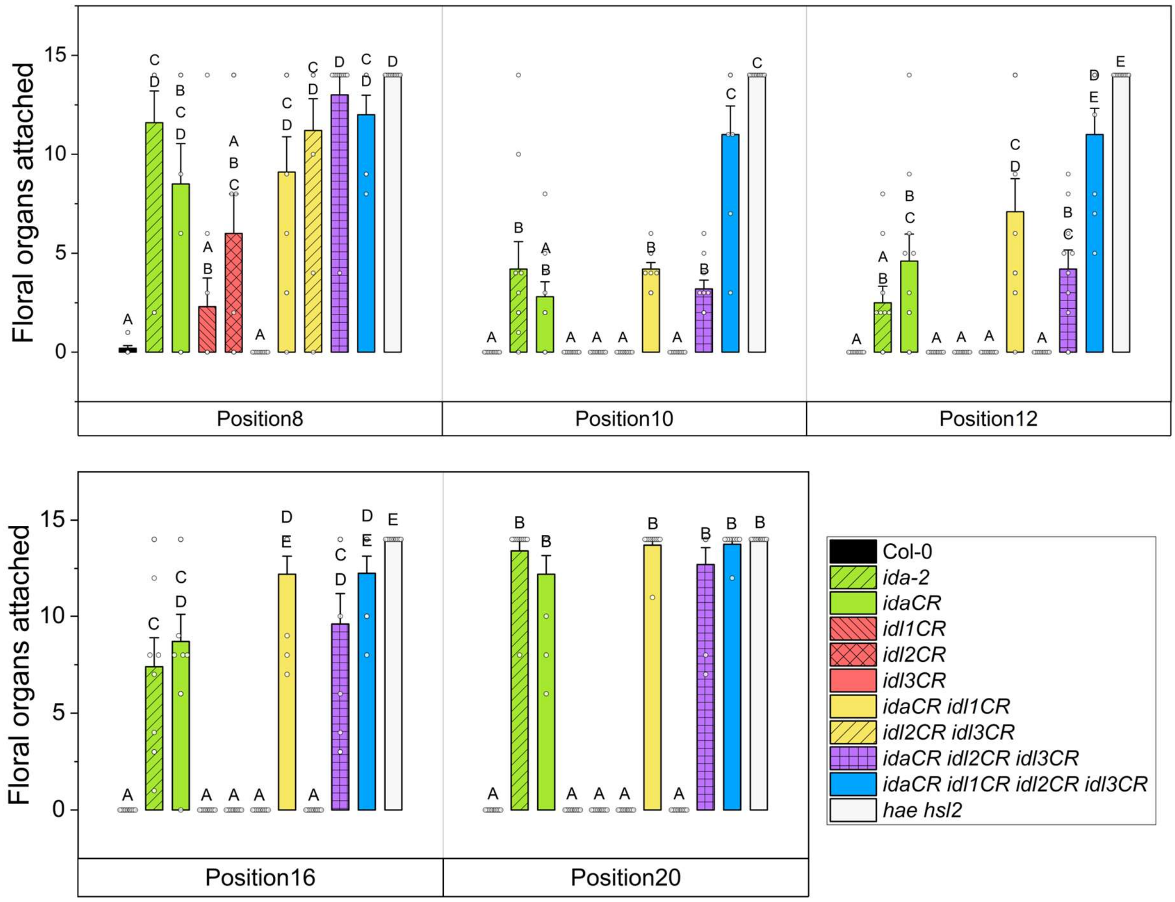


**Supplemental Figure S12.** Quantification of floral organ abscission at positions 8, 10, 12, 16 and 20. This figure contains an extended dataset to the one shown in the main Figure 5F. Letters denote groups with statistically significant means in one way ANOVA and post hoc Bonferroni pairwise comparisons. Statistics shown here were calculated independently at each floral position. Bars show the mean, and whiskers the SEM. Datapoints are overlaid on the barcharts and each represent the number of floral organs attached for each plant phenotyped.
